# A resource-based mechanistic framework for castration-resistant prostate cancer (CRPC)

**DOI:** 10.1101/2023.09.19.558379

**Authors:** B. Vibishan, B.V. Harshavardhan, Sutirth Dey

## Abstract

Cancer therapy often leads to the selective elimination of drug-sensitive cells from the tumour. This can favour the growth of cells resistant to the therapeutic agent, ultimately causing a tumour relapse. Castration-resistant prostate cancer (CRPC) is a well-characterised instance of this phenomenon. In CRPC, after systemic androgen deprivation therapy (ADT), a subset of drug-resistant cancer cells autonomously produce testosterone, thus enabling tumour regrowth. A previous theoretical study has shown that such a tumour relapse can be delayed by inhibiting the growth of drug-resistant cells using biotic competition from drug-sensitive cells. In this context, the centrality of resource dynamics to intra-tumour competition in the CRPC system indicates clear scope for the construction of theoretical models that can explicitly incorporate the underlying mechanisms of tumour ecology. In the current study, we use a modified logistic framework to model cell-cell interactions in terms of the production and consumption of resources. Our results show that steady state composition of CRPC can be understood as a composite function of the availability and utilisation efficiency of two resources-oxygen and testosterone. In particular, we show that the effect of changing resource availability or use efficiency is conditioned by their general abundance regimes. Testosterone typically functions in trace amounts and thus affects steady state behaviour of the CRPC system differently from oxygen, which is usually available at higher levels. Our data thus indicate that explicit consideration of resource dynamics can produce novel and useful mechanistic understanding of CRPC. Furthermore, such a modelling approach also incorporates variables into the system’s description that can be directly measured in a clinical context. This is therefore a promising avenue of research in cancer ecology that could lead to therapeutic approaches that are more clearly rooted in the biology of CRPC.

**Highlights:**

- Cancer growth and progression can be driven by intra-tumour interactions.
- Prostate cancer cells compete among each other for resources like testosterone and oxygen.
- We model these interactions using a modified logistic model with resource dynamics.
- Equilibrium behaviour can be understood through resource supply and use efficiency.
- Explicitly ecological models could enable better strategies for cancer control.

## 1. Introduction

Prostate cancer is among the leading causes of mortality in men (Cancer Society, 2023), and both its clinical progression and therapy have been the subject of intense theoretical investigation (Basanta et al., 2012, Zhang et al., 2017, Cunningham et al., 2018, West et al., 2020, Zhang et al., 2022). Research over the past few years has revealed an important role for intra-tumor competition in prostate cancer progression, particularly in drug-resistant relapse in advanced castration-resistant prostate cancer (CRPC) (Zhang et al., 2017, Gallaher et al., 2018, Brady-Nicholls et al., 2020). While intra-tumor competition is contingent on a diversity of cancer cell types occupying the tumour microenvironment (Gatenby et al., 2009, Kareva et al., 2015, Carreira et al., 2014, Gedye and Navani, 2022, Madan et al., 2022), such cellular heterogeneity also gains therapeutic relevance when a subset of cancer cells in the tumor are resistant to a given drug treatment. Investigations of such tumour populations comprised of a diversity of drug sensitivity phenotypes have demonstrated that drug-resistant relapse under conventional treatment approaches is primarily due to selective elimination of drug-sensitive cells from the tumour by therapy, which in turn allows drug-resistant cells to proliferate more freely (Gatenby et al., 2009, Marusyk et al., 2014, Sahoo et al., 2021, Farrokhian et al., 2022). Current research efforts are therefore aimed at the design of alternative therapeutic regimens that can prevent such competitive release of drug-resistant cells across a variety of cancer types (Hansen and Read, 2020, West et al., 2020).

Prostate cancer is well-characterized in terms of its clinical progression (Gordetsky and Epstein, 2016, Montironi et al., 2018) and the emergence of advanced castration-resistant prostate cancer (CRPC) is considered to be driven primarily by cancer cell types that are resistant to chemical suppression of systemic androgen growth factors through androgen deprivation therapy (ADT) (Culig et al., 1999, Grasso et al., 2012, Fontana et al., 2022). In this context, progress towards alternative strategies to treat CRPC that can better manage drug resistance has been driven to a great degree by a model by Cunningham et al. (Cunningham et al., 2018), which used a logistic framework to cast CRPC as a system of three cell types in density-dependent competition with each other. Uniquely for CRPC, one of these cell types can drive tumour expansion by autonomous production of the androgen, testosterone, which two of the three tumour cell types require for their growth. This adds a commensal dimension to the existing competitive interactions in the system between producers and consumers of testosterone. Cell-autonomous production of testosterone is also thought to drive the clinical progression of prostate cancer towards castration resistance by enabling tumour growth without systemic testosterone supply, which, as mentioned earlier, is directly relevant to the development of CRPC following androgen-deprivation therapy (ADT).

Cunningham et al. (Cunningham et al., 2018) found that the interaction strengths between the cancer cell types, as described by the interaction coefficients, could be used to classify the dynamics and steady states of the model as “Best Responders”, “Responders” or “Non-responders”, to a drug that inhibits testosterone production. Only the testosterone-dependent cell types are sensitive to the inhibitory drug. Therefore, in absence of inter-conversion between these cell types through mutation, a higher frequency of the testosterone-independent cell type will lead to an increasingly drug-resistant tumour population. The relative frequencies of testosterone-dependent cell types therefore determines the “responsiveness” category to which the tumour population is assigned. As with any logistic framework, the steady state stability and cell-type composition in such a model are determined by the values of the interaction coefficients. Thus, the stratification of model behaviour based on drug responsiveness is essentially the identification of qualitatively different points in the parameter space for interaction coefficients.

The Cunningham et al. model represents the use of a classic ecological model of interspecific competition to functionalise tumour heterogeneity within a framework for therapeutic intervention in CRPC. While models of this kind provide valuable phenomenological insights into the progression of CRPC, it is difficult to relate their findings to physiological processes operating within a tumour community. Parameters like the interaction coefficients are usually not directly informative about the underlying mechanisms of cell-cell interactions, and the lack of direct mechanistic links to tumour processes could limit the translational applicability of such models.

Classical theoretical ecology, on the other hand, offers other conceptual frameworks that more explicitly account for the mechanisms of inter-specific interactions. Resource-consumer models represent one such framework within theoretical ecology that cast interspecific interactions entirely in terms of production, supply and consumption of, and competition for, physically explicit resources (Tilman, 1980, Grover, 1997, Muscarella and O’Dwyer, 2020). Their mechanistic roots arguably makes these models harder to parameterize than a logistic framework, which might explain their limited use in cancer systems so far (Kareva et al., 2015). Nevertheless, a resource dynamics based approach offers considerable potential to relate tumour ecology to measurable quantities within the tumour microenvironment.

In this study, we add a mechanistic element to the phenomenological description of the Cunningham et al. model by including the temporal dynamics of two resources-oxygen and testosterone. Oxygen, or the lack thereof (i.e. hypoxia), has been implicated in at least three separate cancer hallmarks including deregulated cellular energetics, angiogenesis, and metastasis (Hanahan and Weinberg, 2011), which makes it a biologically-pertinent inclusion in this model. As recognized by earlier work, testosterone is indispensable to any mechanistic description of the prostate tissue, cancerous or otherwise, as androgens are an integral part of prostate homeostasis (Mohler et al., 2004, Page et al., 2006, Calistro Alvarado, 2010). Furthermore, the demonstrated relevance of within-tumor testosterone production for the progression of advanced prostate cancer (Titus et al., 2005, Montgomery et al., 2008, Stanbrough et al., 2006, Watson et al., 2015) makes it a germane addition to the model.

The inclusion of resource dynamics in our model entails additional processes like cellular resource production and consumption that must be parameterised, and wherever possible, we have determined these parameter values based on available empirical measurements of the corresponding processes. Within this theoretical framework, our results enable us to construct an understanding of the steady state behaviour of the CRPC system in terms of the differential resource use properties of three distinct types of prostate cancer cells. Our data broadly demonstrate that resource consumption processes could potentially elucidate underlying mechanistic links within a system that is otherwise largely phenomenological. In particular, we find that the efficiency of resource use and abundance regimes of resources both play important roles in determining the steady-state composition of the tumour. We also identify resource use efficiency as an interesting aspect of tumour-intrinsic properties that could potentially be used to predict the sensitivity and responsiveness of a given tumour to testosterone-targeting therapy.

## 2 Methods

### 2.1 Model framework

Figure 1 shows a schematic with the main components of the current model. Following the Cunningham et al. model, we use three separate logistic equations (Equation 1) with density-dependent competition to describe the dynamics of each of the three types of cancer cells: *T* ^+^, which requires testosterone for growth but cannot synthesise it; *T*^*p*^, which also requires testosterone for growth but is capable of autonomous testosterone production; and *T*^*−*^, which neither requires nor produces testosterone. We use two ODEs to describe the dynamics of two resources-oxygen (Equation 2) and testosterone (Equation 3)-that contribute directly to cell growth (Figure 1A).

**Figure 1.**
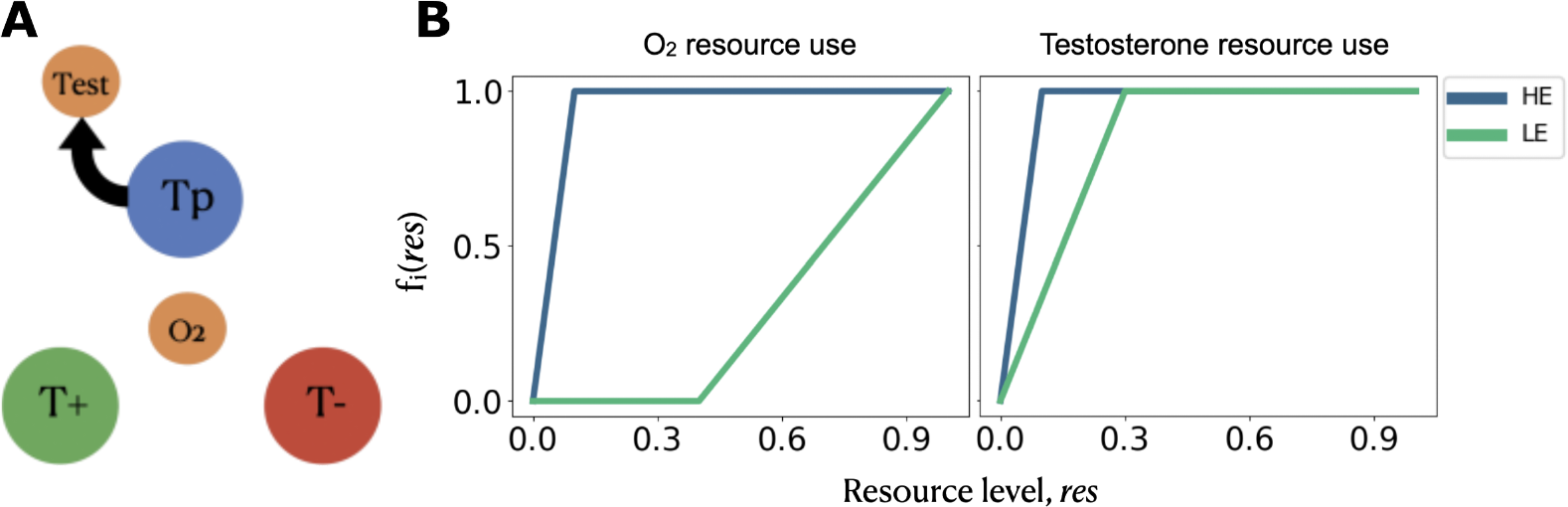
Schematic showing the components of the current model. (A) Three cell types of CRPC, with oxygen as the common resource and testosterone (Test) as the -produced resource. (B) Functional forms used to describe the two resource use efficiencies. HE-high use efficiency, LE-low use efficiency. See main text for descriptions of the cell types and resource use efficiencies.

In order to connect resource levels to cell abundances, we model the cell type-specific carrying capacities as dynamic functions of real-time resource levels according to their respective consumption modalities detailed further below, such that competitive interactions in the model are realised through the levels of each resource. The complete set of equations are as follows:

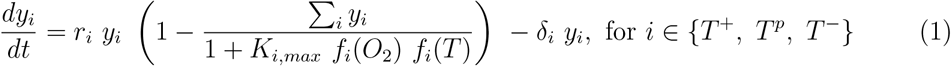

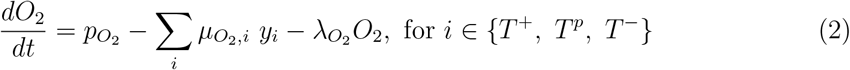

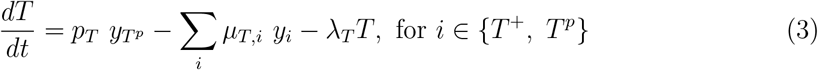

where *r*_*i*_, *y*_*i*_ and *δ*_*i*_ are the growth rate, abundance and death rate for cell type *i*, and *p*_*j*_, *µ*_*j, i*_ and *λ*_*j, i*_ are the production, consumption and decay rates for cell type *i* and resource *j*. We model oxygen as a common resource that is supplied externally in a cell-independent manner reflecting baseline blood supply. It is consumed by all three cell types and leaks out of the tumour tissue at some constant rate. Since castration-resistant prostate cancer arises after some period of androgen deprivation therapy (Mohler et al., 2004), we initially assume that there is no external supply of testosterone, such that any testosterone in the system is derived solely from production by *T*^*p*^. As explained below, this is an assumption that we eventually relax to consider CRPC with external testosterone supply. Since *T* ^*−*^ is testosterone-independent, testosterone consumption is only due to *T*^*p*^ and *T* ^+^, in addition to some constant leakage out of the tumour as with oxygen.

*K*_*i,max*_ is the maximum carrying capacity for each cell type and this is further tuned by the availability of resources through the response function, *f*_*i*_(*res*). As a heuristic based on other models (Ghaffarizadeh et al., 2018), we assume *f*_*i*_ to be a piecewise linear function valued in [0, 1], such that it remains at zero for resource levels below a certain lower threshold value. Once the lower threshold is crossed, *f*_*i*_ increases linearly up to one at some upper threshold value, and stays at one thereafter. Figure 1B shows the general shape of *f*_*i*_ for both resources and this is discussed further in the following section, but we note here that this form allows the effective carrying capacity, as represented by the whole of the denominator in Equation 1, to respond dynamically to the amount of resources available in the environment. When *f*_*i*_(*res*) = 0, the denominator is 1, and for abundant resource availability such that *f*_*i*_(*res*) = 1, the denominator approaches *K*_*i,max*_.

### 2.2 Parameterisation and exploration

Table 1 gives a complete list of all the parameter values and their ranges used in the model. As noted above, we have tried to derive as many of these from empirical sources as possible, such that our model is rooted as far as possible in the biology of the system.

**Table 1:**
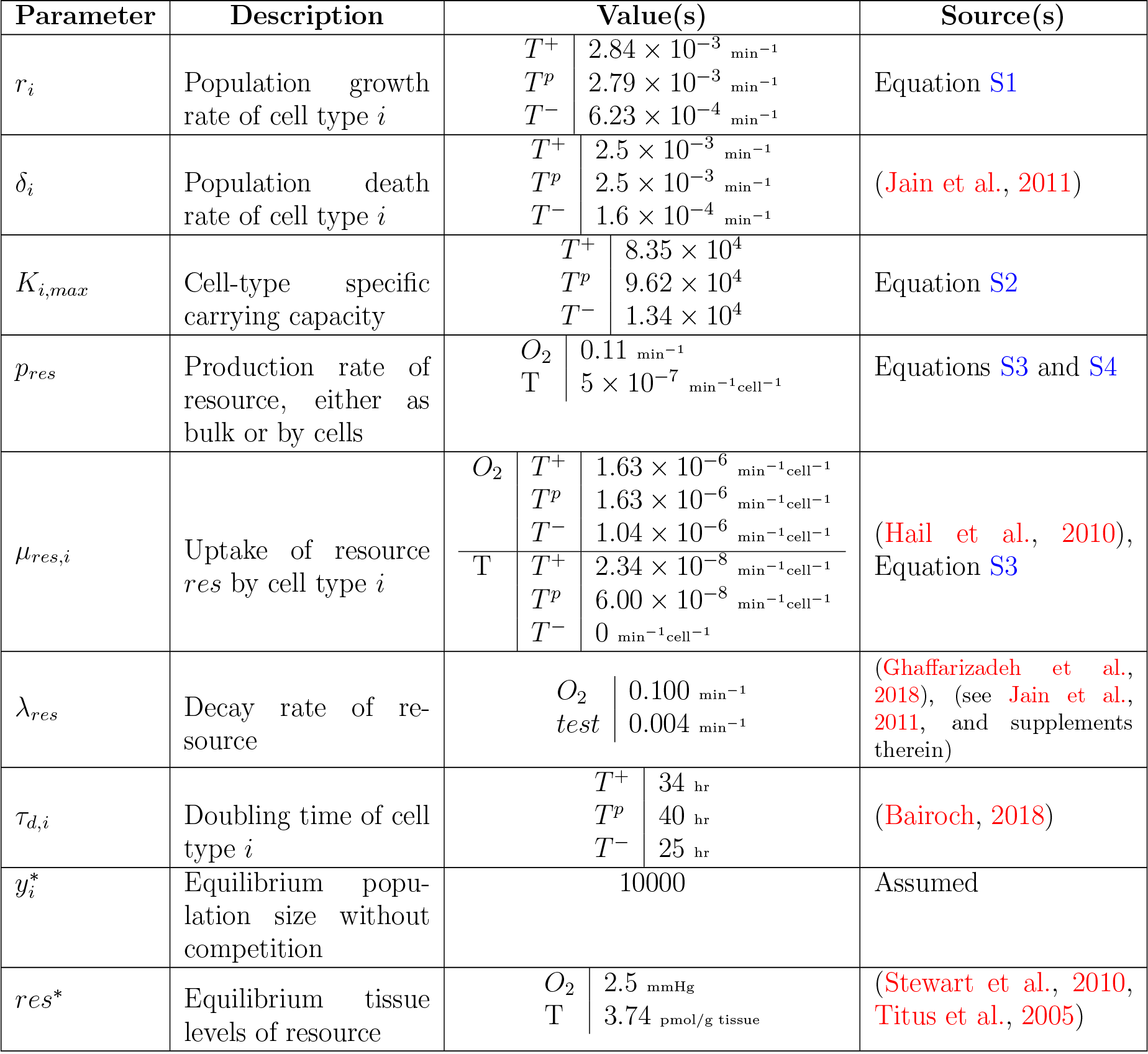
List of all parameters used in the current model. T stands for testosterone. Units of all resource-related parameters do not include dimensions of absolute abundance or concentration as these are normalised against *res*^*\**^, and absolute values can be obtained by multiplying the parameter by *res*^*\**^. Cell-type specific values of *K*_*i,max*_ correspond to the value for which an effective carrying capacity of 10000 is realised for that cell type growing without other competitors and abundant resource availability. Details of normalisation of resource parameters and derivation of growth parameters are given in the Supplementary Text.

Growth and death rates for the cell types are in units of min^*−*1^ and derived from the doubling times of corresponding cell lines (see Table 1 and Supplementary Text). As the growth and death rates are different between the cell types, we derive *K*_*i,max*_ for each cell type grown individually without resource limitations (i.e. *f*_*i*_(*T*) = *f*_*i*_(*O*_2_) = 1) to achieve the same effective carrying capacity of 10000 (Equation S2). We rescale all resource levels such that a resource level of corresponds to the empirically measured steady state value of that resource in prostate cancer tissue. This enables us to sidestep a detailed treatment of the stoichiometry of two very different resources. Setting Equation 2 to zero, we rescale empirically-derived cell-type specific consumption rates and decay rate for oxygen, and use these to calculate the steady state supply rate for oxygen assuming a population of only *T* ^*−*^ cells. Similarly, setting Equation 3 to zero, we use rescaled values of the production rate and natural decay rates for testosterone to calculate testosterone consumption rate for *T*^*p*^. This rate is then used to determine the testosterone consumption rate for *T* ^+^. We restrict all equations to a biologically plausible space by requiring that 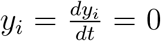, *if y*_*i*_ *<* 1 and only allowing non-zero population and resource abundances. Details of the parameterisation and normalization are given in the Supplementary Text.

The efficiency of resource conversion into cellular growth is an important aspect of any resource-based conception of growth and competition. Resource efficiency is equally relevant to cancer, simply due to the diversity of niches and corresponding metabolic phenotypes within any solid tumour (Zheng, 2012, Birsoy et al., 2014, Loponte et al., 2019). Since environmental heterogeneity is well-documented in solid tumors, the availability and efficiency of use of oxygen could vary substantially between different regions of a tumour (Zheng, 2012). Similarly, variation has been identified across prostate cancer cell lines from several sources in their response to the availability and amount of testosterone in the system (Gregory et al., 2001, Chuu et al., 2011). Resource use efficiency could therefore underlie distinct metabolic strategies within a heterogeneous tumour population. We consider two distinct kinds of such metabolic strategies in our model for each resource, based on the use efficiency of each resource. We choose two distinct forms of the function *f*_*i*_, corresponding to how much resource is required to realise the maximum carrying capacity in Equation 2. The *f*_*i*_ for high use efficiency (Figure 1B, blue line) reaches 1 at a lower resource level than that for low efficiency (Figure 1B, green line), thus requiring a lesser amount of resource to realize maximum carrying capacity.

In addition to resource use, we also explore the effect of initial population size on the steady state composition of the system. This stems from the expectation that as the sole producer of testosterone, absolute abundance of *T*^*p*^ could have a significant impact on the balance of resource-based competition in the system.

All ODEs were simulated numerically using the LSODA algorithm provided by the `scipy.integrate.ode` function in Python, until a plateau was observed. Initial data suggested that a plateau was reached within *∼* 1000 days of model time for most parameter combinations. We therefore ran all simulations for 1000 days initially and subsequently extended the simulation time for those cases where a steady state had not been reached. Supplementary Figures S2-S5 show the time series for all the steady states examined in the main text along with the final run time in each case. All simulations reached steady state within 2500 days, except for only one combination of low oxygen use efficiency and high testosterone use efficiency, and initial population size of 500, which was simulated for 5000 days. Custom Python scripts have been used for all the simulations, data analysis and plotting.

## 3 Results

Our model aims to investigate intra-tumour cell dynamics in CRPC in terms of the availability of and competition for resources. Specifically, we study the effects of changing resource availability under varying conditions of resource use efficiency and population size. Considering two distinct levels of supply for both oxygen and testosterone, we define four resource supply states under which we examine steady state tumour composition. The first state is when **neither** resource is supplemented externally, testosterone production is entirely from *T*^*p*^, and oxygen supply is at a low level, as in a hypoxic tumour. In the second state, **only oxygen** is supplemented and we use a higher value of the oxygen supply rate, while testosterone production remains restricted to *T*^*p*^. In the third state, there is **only testosterone** supplementation, implemented by the inclusion of a constant supply term independent of *T*^*p*^ in Equation 3, while oxygen supply rate is low. In the fourth state, **both** resources are supplemented, oxygen supply rate is high, and external supply of testosterone is included.

We evaluate model outcomes based on whether or not *T*^*p*^ and *T* ^+^ are able to persist at steady state. In the context of therapy, we consider the competitive exclusion of the testosterone-dependent cell types by *T*^*−*^ an unfavourable outcome, the reasons for which will be discussed later. We classify model behaviour into four distinct tumour responses based on the particular resource supply state in which steady-state persistence of *T*^*p*^ and *T* ^+^ occurs. The first is a T-type response, which signifies that external supplementation of testosterone is sufficient for *T*^*p*^ *− T* ^+^ persistence at steady state. The second is an O-type response, which signifies that external supplementation of *O*_2_ alone is sufficient for *T*^*p*^ *− T* ^+^ persistence. The third is a TO-type response signifying that external supplementation of both resources is required for *T*^*p*^ *− T* ^+^ persistence. The fourth response is called an N-type one, and indicates that *T*^*p*^ *− T* ^+^ persistence is not possible under any of the resource supply states tested here. Unless otherwise stated, all simulations are initialized with equal numbers of all three cell types and resource use efficiencies are identical across all three cell types. In the following sections, we illustrate how both resource use efficiency and supply states can play key roles in determining the overall tumour response as defined above.

### 3.1 Steady state composition under high resource use efficiency: a mix of tumour responses

Figure 2 shows the steady-state composition of the tumour across the four resource supply states when both resources are used at the same efficiency, either both high (Figure 2A) or both low (Figure 2B). At high use efficiency (Figure 2A), for any initial population size, *T*^*p*^ *− T* ^+^ are competitively excluded by *T* ^*−*^ when neither resource is supplemented externally.

**Figure 2.**
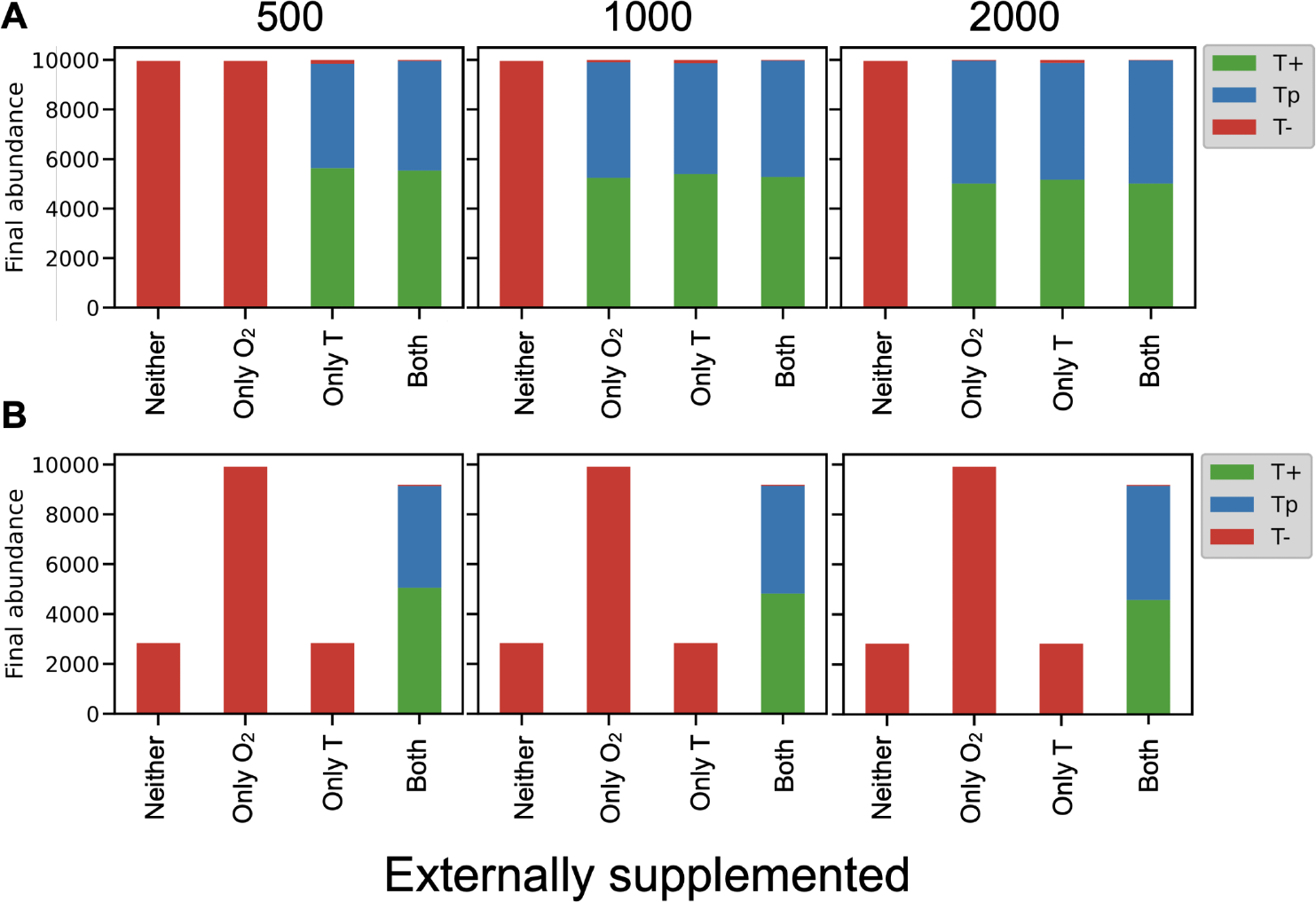
Steady state behaviour under identical use efficiencies of both resources. Abundances of all three cell types, under (A) high use efficiency of both resources, and (B) low use efficiency of both resources; initial population sizes are given on top of each subpanel and equal numbers of each cell type were seeded in the beginning. Without external supplementation, *p*_*O*2_ = 0.06 min^*−*1^, *O*_2_ at *t* = 0 was set to 0, and testosterone was produced by *T*^*p*^ at a rate *p*_*T*_ = 5 *×* 10^*−*7^ min^*−*1^ per cell. Under external supplementation, *p*_*O*2_ = 0.11 min^*−*1^, *O*_2_ at *t* = 0 was set to 0.5 and an additional supply of 0.001 min^*−*1^ was added to Equation 3. *T* at *t* = 0 was always 0. High use efficiency (i.e. panel A) usually leads to either a T- or T/O-type tumour response depending on the initial population size. Low use efficiency (i.e. panel B) produces an obligate TO-type tumour response regardless of initial population size. Further elaboration on tumour response types are in the main text.

This is intuitive as *T*^*p*^ and *T* ^+^ are dependent on both testosterone and oxygen for growth, and lack of external supplementation of both resources puts them at a competitive disadvantage relative to *T* ^*−*^ which are limited only by oxygen. At population size 500, external supply of testosterone is sufficient to ensure *T*^*p*^ *− T* ^+^ persistence, indicating a T-type response. However, for higher initial population sizes, external supply of either or both resources can ensure *T*^*p*^ *− T* ^+^ persistence, suggesting a T- and O-type tumour response. Both these responses can be understood by identifying the limiting resource in each case. In the absence of external testosterone supply, *T*^*p*^ cells are the only source of testosterone in the system. A small initial population with few *T*^*p*^ cells and no external testosterone supply is therefore limited by testosterone, leading to a T-type tumour response. Larger initial population sizes also entail higher *T*^*p*^ frequency, thus relieving testosterone limitation. Under these circumstances, external supply of either one of the resources is sufficient to support *T*^*p*^ *− T* ^+^ growth and steady-state persistence. Taken together, our data indicate that high resource use efficiency generally favours *T*^*p*^ *− T* ^+^ persistence at steady state under a range of resource supply conditions, although this tendency is affected by the initial population size.

Under low use efficiency of both resources, Figure 2B shows that supply of both resources is necessary for *T*^*p*^ *− T* ^+^ persistence at steady state, indicating a TO-type tumour response. Contrary to high efficiency (Figure 2A), the TO-type response under low use efficiency is unaffected by the initial population size. We also observe that under low efficiency, low supply of oxygen leads to a smaller steady-state population size while the lack of external testosterone supply does not have such an effect on the steady-state population size. This disparity is explicable based on the functional form of low use efficiency of oxygen, which allows only a fraction of the maximum carrying capacity to be realised under the oxygen-limiting conditions imposed by limited external supply. It is also worth noting that since testosterone use efficiency is also low, *T*^*p*^ *− T* ^+^ growth is limited strongly by both oxygen and testosterone at all times, and therefore requires external supplementation of both resources for their persistence. This leads to a TO-type response.

### 3.2 Mismatched resource use efficiencies reveal the lability of tumour responses

So far, we have only dealt with identical use efficiency of both resources-either both high or both low. We now consider responses to resource supplies of tumors with one resource used at high efficiency and the other at low efficiency. As Figure 3 (left) shows, low testosterone use efficiency is the simpler of the two in terms of tumour responses, as it leads to an obligate T-type response. This implies that *T*^*p*^ *− T* ^+^ persistence requires external testosterone supply regardless of the initial population size, and as Figures 3A, C and E make clear, testosterone is both necessary and sufficient for *T*^*p*^ *− T* ^+^ persistence. As observed earlier, lower testosterone abundance does not affect the steady state population size.

**Figure 3.**
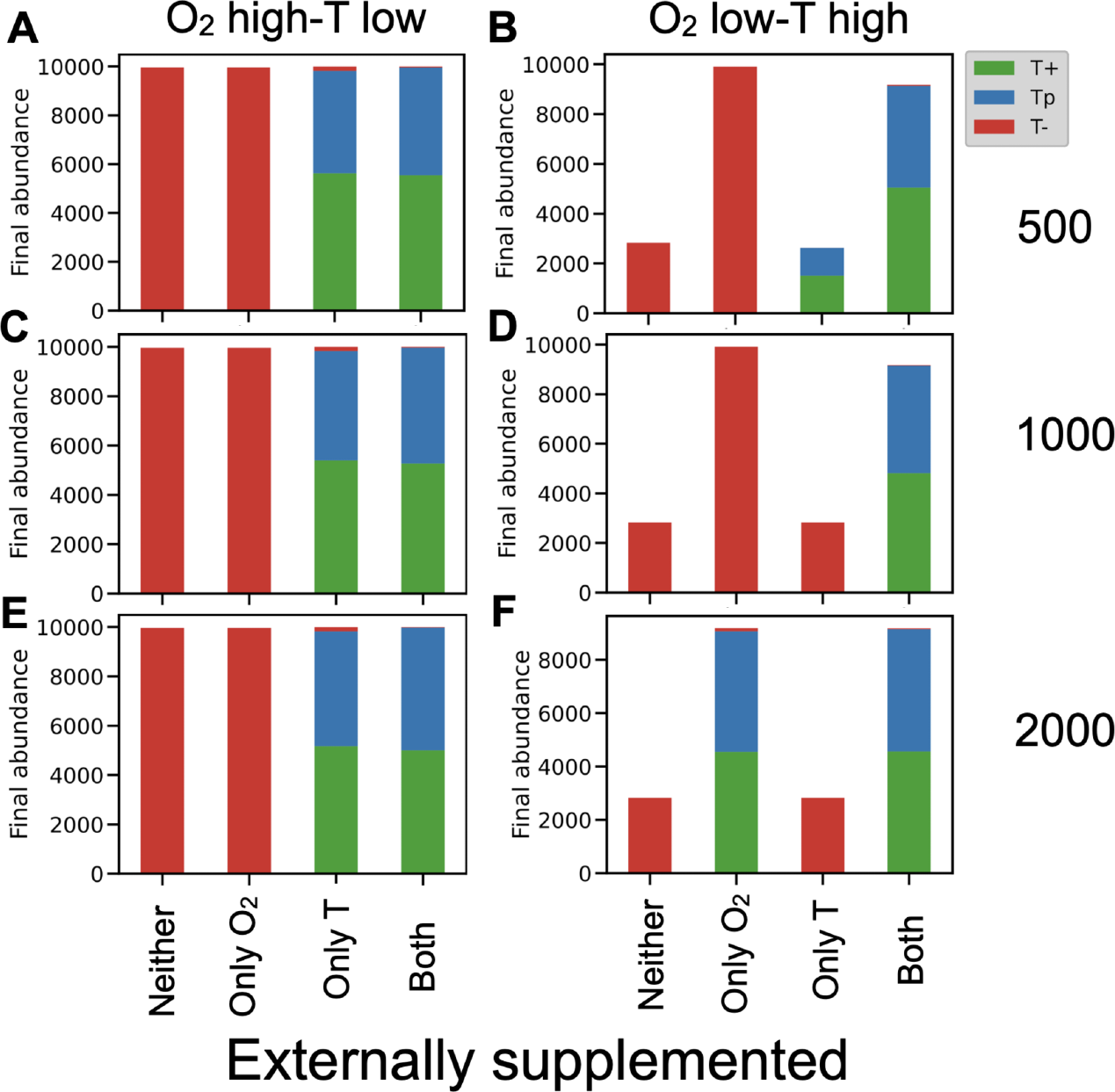
Steady state behaviour under mismatched resource use efficiencies. Abundances of all three cell for (A, C, E) low use efficiency of testosterone only and (B, D, F) low use efficiency of oxygen only; initial population sizes vary across rows as given on the right end of each row, and equal numbers of each cell type were seeded in the beginning. Resource supplies were parameterised as mentioned earlier. T stands for testosterone. When testosterone use efficiency is low (A, C, E), tumour responses are obligately T-type regardless of initial population size. On the other hand, low use efficiency of oxygen (B, D, F) leads to shifting tumour responses, from T-type for small initial population size (B), to TO-type response (D), to an O-type response (F). See main text for further explanation.

Low use efficiency of oxygen coupled with high testosterone use efficiency presents a more complicated picture. As shown in Figure 3 (panels B, D and F), steady state population size responds to low oxygen abundance as expected, leading to a smaller final population. However, when oxygen and/or testosterone are supplied, tumour responses in terms of steady state composition show a greater degree of variability. Following the trend across Figure 3B, D and F from top to bottom, at small initial population size (Figure 3B), the lower frequency of *T*^*p*^ cells leads to testosterone limitation and the tumour shows a T-type response. Given that *T*^*p*^*−T* ^+^ are already limited by testosterone, external supply of oxygen under small initial population size favours *T* ^*−*^. Intermediate initial population size increases the *T*^*p*^ frequency, but in the absence of external testosterone supply, testosterone production by *T*^*p*^ alone is insufficient for *T*^*p*^ *− T* ^+^ persistence. Since oxygen is already limiting, this leads to a TO-type response overall (Figure 3D). Further increase in the initial population size increases *T*^*p*^ frequency, and this increase ameliorates the initial testosterone limitation. However, higher total cell number under low oxygen use efficiency makes oxygen the limiting resource in turn, and this leads to an O-type response (Figure 3F). Low use efficiency of oxygen therefore reveals the potential for population size to determine which resource is actually limiting and how tumour populations respond to resource supply states.

### 3.3 Oxygen supply rate could affect producer-consumer cell abundance balance

Figure 4 shows part of a finer exploration of the resource-supply parameter space, in which we vary the supply rates of both resources over a range of values. Under low use efficiency of only oxygen (Figure 4A), we find that increasing the supply rate of oxygen increases the proportion of *T* ^+^ at steady state relative to *T*^*p*^ (*cf* Figure 4A and B). Low efficiency use of testosterone, on the other hand, limits the viable resource supply region, such that population growth occurs for only the highest testosterone supply rate when its use efficiency is low (Figure 4B; note that the x-axis here has only the highest value of testosterone supply rate).

**Figure 4.**
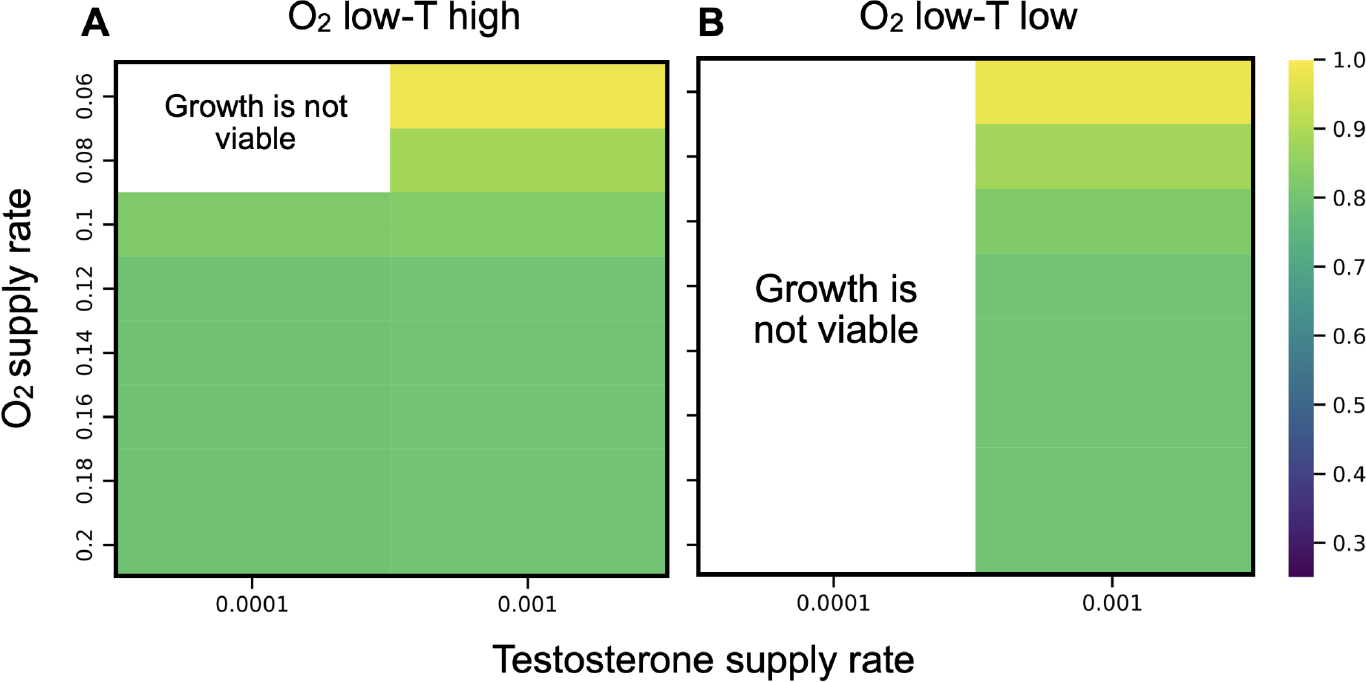
Producer-consumer balance with varying oxygen supply. Steady state ratio of *T*^*p*^ against *T* ^+^ across a range of supply rates of oxygen and testosterone for low use efficiency of oxygen and either (A) high use efficiency or (B) low use efficiency of testosterone. Note that lower supply rates of either resource, below 0.0001 min^*−*1^ for testosterone and below 0.06 min^*−*1^ for oxygen, could not support any cell growth at all and are therefore not shown here. Empty spaces marked as “Growth is not viable” are resource supply states for which *T*^*p*^ and *T* ^+^ both go extinct. Oxygen supply rates on the y-axis increase from top to bottom. In all cases, simulations were initialised with equal numbers of all three cell types, with an initial population size of 500 and run for 2500 days.

It is also worth noting that in the supply cases shown here, the steady-state population is almost entirely comprised of *T*^*p*^ and *T* ^+^, indicating that under sufficient resource abundance, *T* ^*−*^ is competitively excluded. As further elucidated in Supplementary Figure S6, below some lower threshold supply rates of oxygen and testosterone, *T*^*p*^ *− T* ^+^ are both driven to extinction at steady state. While the data are not shown here, we also find that below some other threshold supply of oxygen lower still, all three cell types are driven to extinction. Testosterone supply and its use efficiency therefore seem to result in binary outcomes in terms of steady-state *T*^*p*^ *− T* ^+^ persistence. On the other hand, under conditions when testosterone availability is sufficient for -persistence, higher oxygen availability favours a greater proportion of *T* ^+^ at steady state.

### 3.4 Oxygen and testosterone function at different abundance regimes

The previous sections show that across resource efficiencies, oxygen and testosterone have qualitatively different effects on steady state population sizes and tumour compositions. To investigate these differences in further detail, we show part of the parameter exploration for the choice of upper and lower thresholds for *f*_*i*_(*O*_2_) and *f*_*i*_(*T*) in Figure 5A-B, for a population of *T*^*p*^ cells alone. The heat maps clearly demonstrate that when testosterone is only produced by *T*^*p*^ and oxygen is supplied externally, *T*^*p*^ growth is viable across the whole parameter space of upper and lower thresholds for oxygen (Figure 5A). However, *T*^*p*^ goes extinct for testosterone lower thresholds of 0.2 or greater (Figure 5B). Comparing this with a representative simulated time series of resource levels in Figure 5C, we see that in the initial few days, testosterone levels remain slightly below 0.2 whereas oxygen levels increase rapidly to stabilize close to one. This distinction in turn stems from the fact that oxygen and testosterone are produced and consumed at vastly different rates in the system, leading to different abundances at steady state as well as different dynamics towards the steady state.

**Figure 5.**
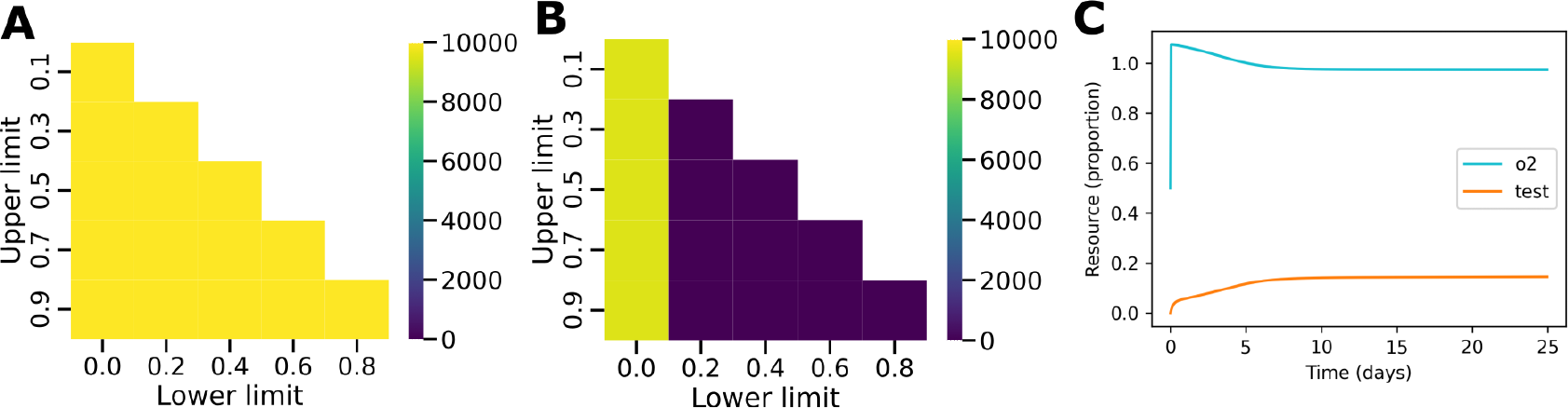
Abundance regimes of oxygen vs testosterone. Steady state abundances of a population of *T*^*p*^ cells grown without resource limitation and in the absence of other cell types, under a range of thresholds for *f*_*i*_ of (A) oxygen and (B) testosterone, and (C) representative time series of resource dynamics in the first 25 days. For panels (A) and (B), upper and lower threshold values correspond to the upper and lower inflection points of *f*_*i*_ as shown in Figure 1B. The missing upper diagonal values of (A) and (B) are not valid threshold values of *f*_*i*_ as the lower limit must be less than the upper limit by definition. For (C), equal numbers of all three cell types were initialised, initial total population size was 2000, lower and upper thresholds were 0 and 1 respectively for *f*_*i*_(*O*_2_), 0 and 0.1 for 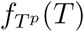, and 0 and 0.05 for *f*_*T*_ + (*T*), and 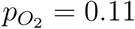 min^*−*1^.

Figure 5 altogether demonstrates that the difference in steady state resource abundance corresponds to the parameter space of *f*_*i*_(*O*_2_) and *f*_*i*_(*T*) where cell growth is viable. The extent to which the upper and lower thresholds can be varied for *f*_*i*_(*T*) is therefore much more restricted than for *f*_*i*_(*O*_2_), primarily due to differences in the underlying resource dynamics.

## 4 Discussion

The current study goes beyond previous investigations (Cunningham et al., 2018) by allowing us to describe system behaviour in terms of the availability and consumption of resources that are pertinent to the biology of CRPC. In the absence of external supplementation, oxygen in our model is usually present at much higher levels than testosterone (Figure 5), which is consistent with measurements from the tumour microenvironment (Dillard et al., 2008, Stewart et al., 2010). As *T*^*p*^ and *T* ^+^ are simultaneously limited by two resources, they are driven to extinction in many parts of the resource efficiency-supply space (Figures 2 and 3). While low resource use efficiency is detrimental to *T*^*p*^ *− T* ^+^ survival overall (Figure 2B), overcoming this through external supply of the limiting resource is context-dependent. Figure 3A shows that when testosterone use efficiency is low, external supply of testosterone is always sufficient to ensure *T*^*p*^ *− T* ^+^ survival. When oxygen use efficiency is low, Figure 3B shows that the initial population size determines which resource is limiting and can therefore support *T*^*p*^ *− T* ^+^ survival if supplied externally. Figure 4 shows a specific part of the resource supply-efficiency parameter in which oxygen supply seemingly favours a higher abundance of *T* ^+^ at steady state. Since *T* ^+^ has a slightly higher doubling rate than *T*^*p*^, it is possible that its growth could be favoured by the presence of higher resource abundance. However, an examination of the corresponding time series (Supplementary Figures S7-S9) suggests that the simulated time window of 2500 days could have been insufficient for the growth curves of *T*^*p*^ and *T* ^+^ to reach a clear plateau in some cases. However, as the time series also indicate, the tumour composition changes so slowly after 2500 days that any effect of supply rates on cell type abundances would be of limited clinical importance.

On the whole, our data suggest that resource use efficiencies could serve as potential tumour-intrinsic properties that could be predictive of tumour composition at steady state. In the case of CRPC, the steady-state composition serves as an indicator of whether the tumour is likely to respond to resource-targeting therapy. As *T*^*p*^ and *T* ^+^ are both testosterone-dependent cell types, their persistence at steady state leads to a tumour that would respond to testosterone inhibition. On the other hand, a higher frequency of *T* ^*−*^ at steady state leads to a tumour that is less responsive to testosterone-targeting therapy as *T* ^*−*^ cells are not dependent on testosterone for growth. Given that the development of personalised cancer therapy is a subject of active research (Martin et al., 2015, Hansen et al., 2017, Marigorta et al., 2017, Gedye and Navani, 2022), our data indicate that the elucidation of resource use strategies in a tumour could form the basis of designing more effective, personalized therapeutic strategies for CRPC cases depending on the extent of resource dependencies within the tumour. We also speculate that the abundance regime in which a resource operates could be an important factor in determining its therapeutic relevance. Testosterone in our model operates at a much lower abundance than oxygen. On one hand, this identifies testosterone as a clear target for therapeutic approaches involving resource supply as it is invariably limiting. However, we also find that model behaviour is highly sensitive to changes in testosterone availability, with very sharp thresholds for steady-state *T*^*p*^ *− T* ^+^ persistence (Supplementary Figure S6). To what extent these thresholds are an artefact of our parameterisation of resource use efficiencies is unclear, and this is a subject of ongoing investigation.

Although we use clinically relevant parameter regimes for our data, our model results have not been fit to longitudinal data from patient samples, unlike some other models in this context (Zhang et al., 2022). Consequently, rather than providing precise quantitative predictions about the progression and dynamics of CRPC, our results point towards the qual-itative patterns and behaviours of such a system. These results therefore could be of value for growing developments in cancer management strategies. As the focus of cancer treatment shifts from complete cure to effective control, identifying and delimiting new ways to achieve such effective control is of vital importance. Cytotoxic chemotherapy weakens intra-tumour competition by depleting the tumour of sensitive cells, but our results suggest that in cases where drug sensitivity is accompanied by unique resource use strategies, sensitive cells could be supported by external supply of the corresponding resource that they use exclusively. This could offer an effective counterweight to chemotherapy by strengthening intra-tumour competition, thus potentially extending the time for which the same therapy can be administered fruitfully. Broadly, these inferences stemming from our model are supported by recent work on a classical resource-consumer model of estrogen and glucose consumption in breast cancer, which used an *R*^*\**^-based conceptual framework to discuss the effects of resource dynamics on intra-tumour competition (Kareva and Brown, 2021). Specifically for prostate cancer, it has been observed that tumours of an LNCaP-derived cell line in athymic mice undergo significant regression and necrosis when the mouse is given a testosterone bolus through an implanted pellet (Umekita et al., 1996). Therapeutic use of testosterone also offers the possibility of improvements in patient-reported Quality of Life indicators that are typically adversely affected by prolonged androgen deprivation (Niraula et al., 2016). The design and optimization of dosing regimens that can use both cytotoxic drugs and externally-supplied resources could therefore be a promising direction for the treatment of CRPC (Nishiyama and Hoshii, 2016, Teply et al., 2018).

It is undoubtedly a huge challenge to reliably pinpoint which resources are limiting and to what extent, given the considerable complexity of the tumour microenvironment. However, even simple identification of tumours in which a resource-based intervention could be effective is a reasonable step forward. For instance, tumours corresponding to high resource efficiencies in our model are likely to be excellent candidates for a resource-based treatment strategy as they are likely to respond to changing resource availabilities without complete loss of the drug-sensitive population. The mapping of our model outcomes to therapeutic feasibility is currently limited both because the model is not fitted to clinical data and because our conceptualization of resource efficiencies is entirely heuristic. That said, our data indicate that a theoretical framework that integrates available information on the biology of resource use in a tumour with its growth dynamics could considerably expand our current toolkit for cancer control. This view has seen increasing advocacy recently (Kareva et al., 2015, Kareva and Brown, 2021) and has particular relevance for endocrine cancers, most of which frequently have a hormone or growth factor signaling axis that is crucial for cancer progression, as exemplified by the importance of EGFR signaling in breast cancer (Osborne et al., 1985, Kareva and Brown, 2021), testosterone signaling in prostate cancer (Mohler et al., 2004), or insulin signaling in pancreatic cancer (Chan et al., 2014, Archetti et al., 2015). More such signaling axes will undoubtedly be identified in the years to come and in each case, tumour composition and size could be seen as a result of the interplay between such signaling and other energetic resources, which offers much scope for further modelling effort.

Realizing the translational value of resource use models in cancer treatment depends on the ability to identify and monitor relevant cellular phenotypes and resource availability. One potentially interesting avenue are the so-called “radiomics” approaches that can integrate quantitative imaging data with additional information about spatial structure, genomics and gene expression to help with diagnosis and monitoring (Gillies et al., 2016). Such analyses have been applied to classify prostate cancer and benign masses accurately, and further separate different kinds of tumours with the same overall Gleason score (3 + 4 vs 4 + 3, for example (Wibmer et al., 2015, Fehr et al., 2015)). The possibility of obtaining such information before and after a therapeutic intervention represents unprecedented scope in the future for precise, patient-driven treatment that can respond to drug-induced ecological and evolutionary changes in a tumour. Theoretical models must therefore be geared to make use of such granular data from the tumour microenvironment to enable an ecologically-informed treatment approach.

## Acknowledgements

The authors acknowledge current and past members, and visiting students of the Population Biology Lab at IISER Pune for extensive critical inputs on previous analyses. The authors also thank Deepak Barua for useful discussions. This project was supported by grant # STR/2021/000021 from Science and Engineering Research Board, Department of Science and Technology (DST), Government of India and internal funding from Indian Institute of Science Education and Research (IISER), Pune. BV was supported by IISER Pune and HBV through the Kishore Vaigyanik Protsahan Yojana (KVPY), DST, Government of India.

## Author contributions

BV and SD conceptualised the study. HBV wrote the original codes for simulations, plotting and analysis, and contributed to the data in Figures 2, 3 and 5. BV contributed additional data and analysis for Figures 2, 3, 4 and 5, and wrote the manuscript with inputs from HBV and SD. All authors approved the final version of the manuscript.

## Supplementary Text

### Derivation of cell growth parameters

For a small initial population and abundant resource supply, Equation 1 can be simplified to an exponential growth process by ignoring the quadratic growth term. Solving this for one population doubling leads to the following expression for the exponential growth rate for each cell type:

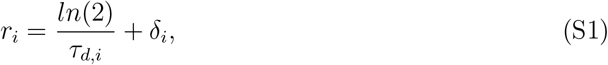

where *r*_*i*_ is the cell-type specific growth rate, *δ*_*i*_ is the cell-type specific death rate as inferred from (Jain et al., 2011), and *τ*_*d,i*_ is the doubling time from available data of cell lines corresponding to each of our cell types (Bairoch, 2018). We take *T* ^+^, *T*^*p*^ and *T* ^*−*^ to correspond to the LNCaP clone FGC (RRID: CVCL 1379), 22Rv1 (RRID: CVCL 1045) and PC-3 (RRID: CVCL 0035) cell lines respectively, and use the doubling times of these cell lines to calculate the growth rates by Equation S1. We assume that the effective carrying capacity for each cell type growing in the absence of other cell types under abundant resource availability to be 10000. Setting the LHS of Equation 1 to zero and rearranging, we obtain the following expression for *K*_*i,max*_:

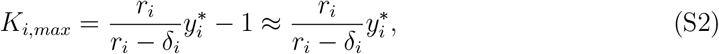

where 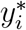 is the steady-state population size of cell type *i ∈ T*^*p*^, *T* ^+^, *T* ^*−*^, growing in the absence of competition from other cell types and abundant resource availability, and 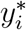 is set to 10000. Substituting this in Equation S2 allows us to calculate cell-type specific values for that would realise a maximum population size of 10000 for each cell type.

### Normalization of resource dynamics

As oxygen and testosterone are produced and consumed at different rates, their stoichiometry is likely to be vastly different. In the interest of simplicity, we neglect considerations of stoichiometry in our model by normalising the abundance of each resource against the empirically measured steady state value of that resource in prostate tissue. Median oxygen tension in prostate cancer tissue is found to be about 2.5 mmHg (Stewart et al., 2010) and we take this to be the steady state level of oxygen in prostate cancer tissue. For a tumour volume of 1mL, this corresponds to a steady state oxygen level of 3.375 nmol. Setting the LHS of Equation 2 to zero, considering a population of only *T* ^*−*^ cells, normalising both *λ*_*O*2_ and *µ*_*O*2,*i*_ against 3.375 nmol, and setting normalised equilibrium oxygen, *O*_2_ to one, we obtain *p*_*O*2_ corresponding to vascularised PCa tissue from Equation S3 as 0.11 units per min. For limited oxygen supply, we assume *p*_*O*2_ = 0.06 units per min.

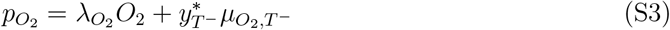

Studies on tumour tissue from patients undergoing ADT find a testosterone level of 3.75 pmol/g tissue in PCa (Titus et al., 2005). As with oxygen, we take this as the steady state value and for a 1mL tumour volume with tissue density 1.03 g/cm^*−*3^ (Stamey et al., 1987), this corresponds to a steady state testosterone level of 3.64 pmol. All rate parameters are divided by this value to give consumption, production and decay rates in units of proportion of steady state resource level. For the rate of testosterone production by *T*^*p*^ cells (*p*_*T*_), empirical sources indicate an absolute rate of about 1100 pg per million cells per 48h (Dillard et al., 2008), which is about 3 *×* 10^*−*10^ units per cell per min after normalization. Initial simulations suggested that this rate of production was too low for to support cell growth from a small initial number of cells within the simulated time. Following a brief parameter space exploration, we assume *p*_*T*_ = 5 *×* 10^*−*7^ units per cell per min to ensure sufficient testosterone availability. Setting Equation 3 to zero, taking normalized equilibrium testosterone, *T* ^*\**^ = 1, and normalizing *λ*_*T*_ to 3.64 pmol, we obtain *µ*_*T,T*_ *p* for a steady-state *T*^*p*^ population of 10000 as 6 *×* 10^*−*8^ units per cell per min from Equation S4 below.

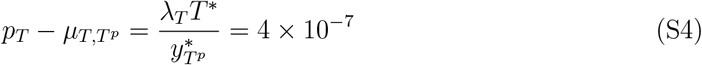

Since direct empirical sources were not available for *T* ^+^ testosterone consumption rate, we explore values over four orders of magnitude from 10^*−*11^ to 10^*−*7^ units per cell per min, with the additional assumption that *T* ^+^ consumed testosterone at a lower rate than that of *T*^*p*^. Supplementary Figure S1 shows the time series of cell growth and resource dynamics for the highest rate of testosterone consumption by *T* ^+^, and we see that *T* ^+^ is outcompeted early in the simulation by *T*^*p*^. Closer examination of the time series data indicates that while both cell types initially decrease in abundance due to resource limitation, *T* ^+^ declines more rapidly and is thus driven to extinction by *T*^*p*^. Since this extinction is independent of the testosterone consumption rate of *T* ^+^, we fix this rate to be a value slightly less than the *T*^*p*^ consumption rate. This is effectively a simplification of the *T*^*p*^ *− T* ^+^ interaction, as it precludes a potential amensal interaction by which *T* ^+^ could consume an excess of the testosterone produced by *T*^*p*^. For external supply of testosterone, we add a constant supply term of 0.001 per min to Equation 3.

## Supplementary Figures

**Supplementary Figure S1:**
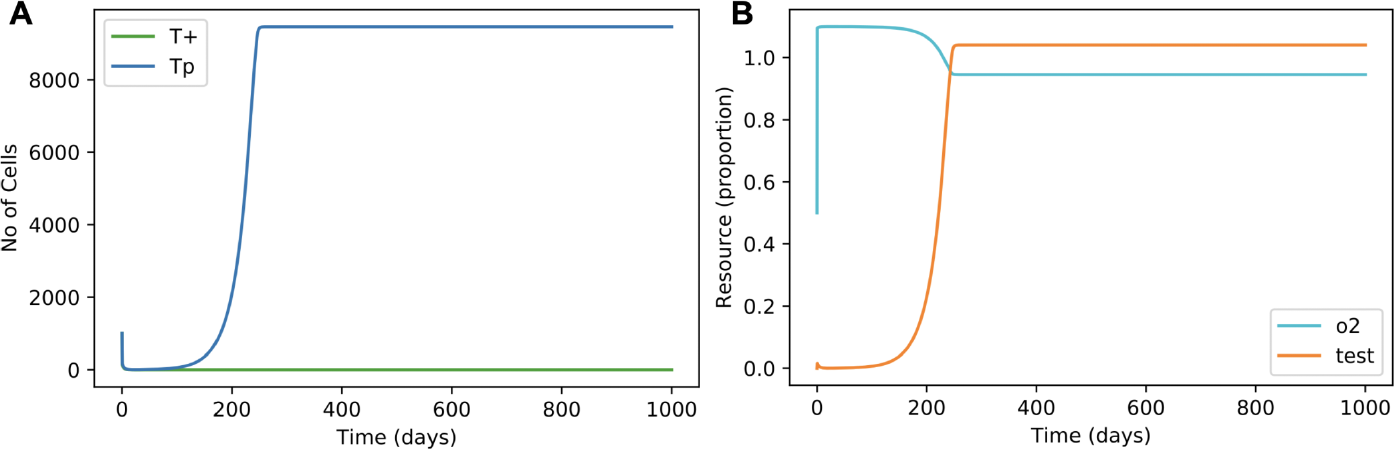
Fixing the testosterone consumption rate for *T* ^+^. Time series of (A) *T*^*p*^ and *T* ^+^ cell growth, and (B) resource dynamics in the absence of *T* ^*−*^. *T*^*p*^ *−T* ^+^ dynamics show that *T* ^+^ is competitively excluded by *T*^*p*^ at steady state. Initial population size is 1000 cells of each type, 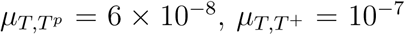, lower limit for the resource use function *f*_*i*_ for both resources and both cell types is zero, and the upper limit is one.

**Supplementary Figure S2:**
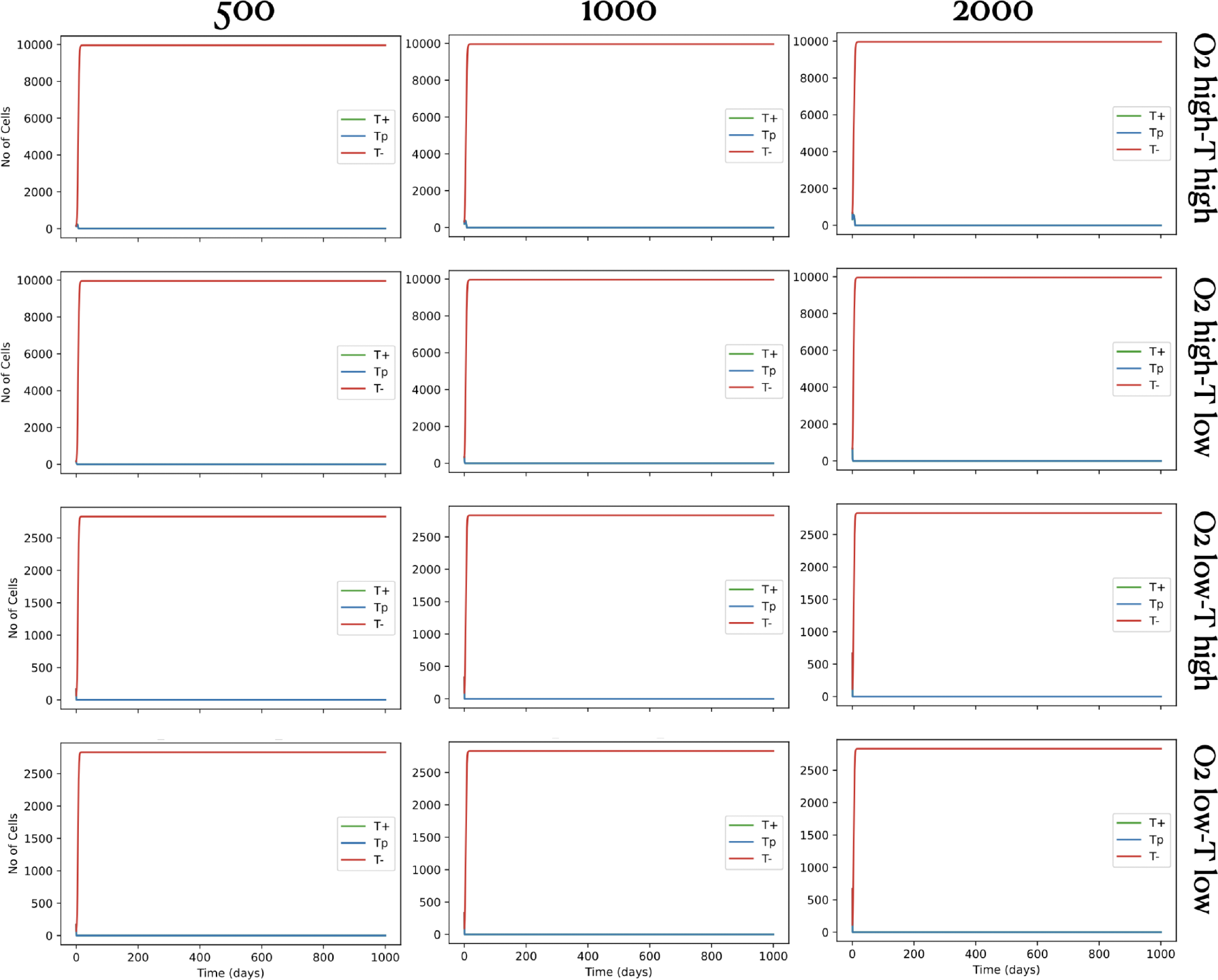
Cell growth time series when <u>neither</u> resource is supplemented externally. Resource use efficiency combinations vary across rows and initial population size across columns. These simulations were run for 1000 days since steady state was seen to be reached within the first few days.

**Supplementary Figure S3:**
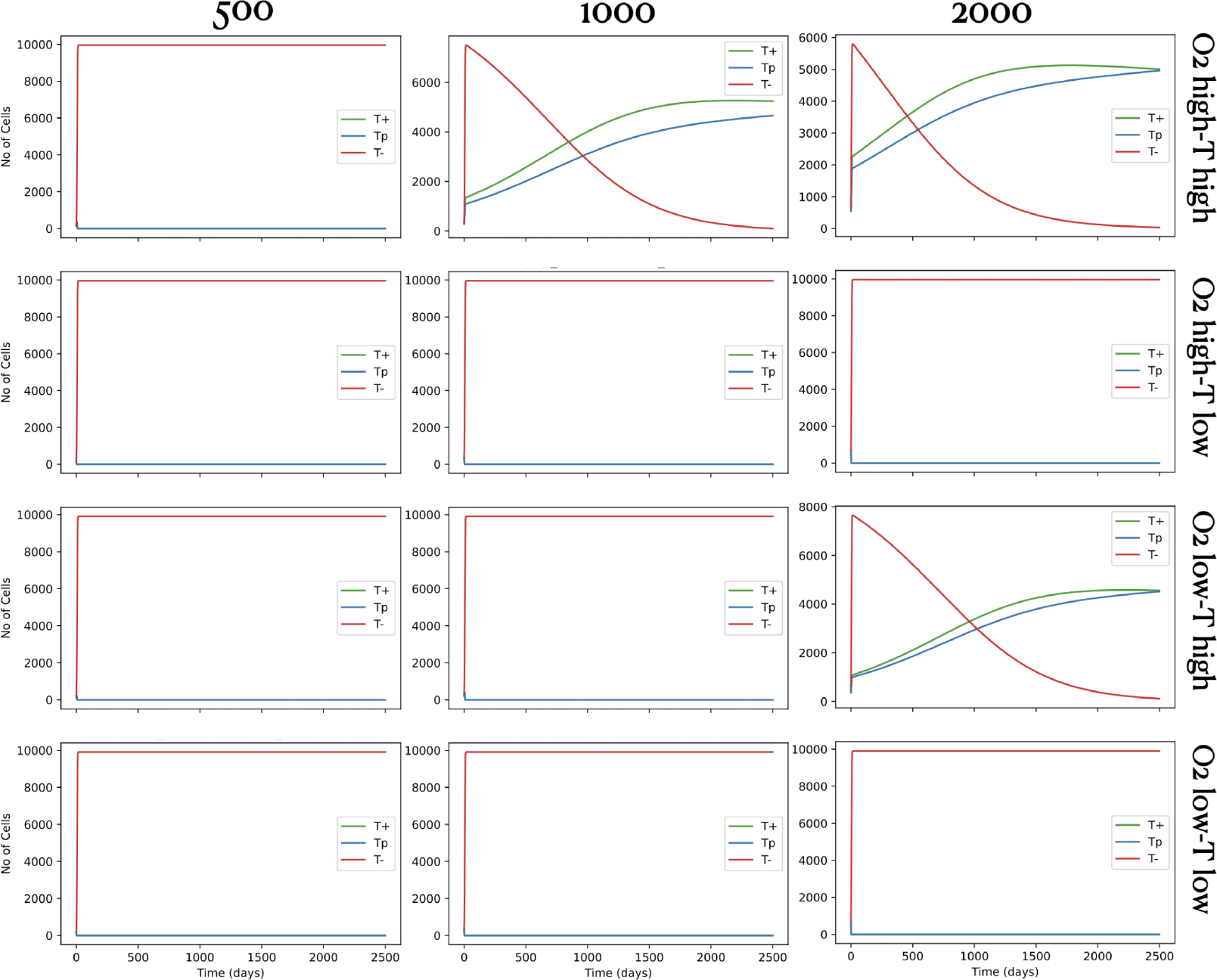
Cell growth time series when <u>only oxygen</u> is supplemented externally. Resource use efficiency combinations vary across rows and initial population size across columns. These simulations were run for 2500 days.

**Supplementary Figure S4:**
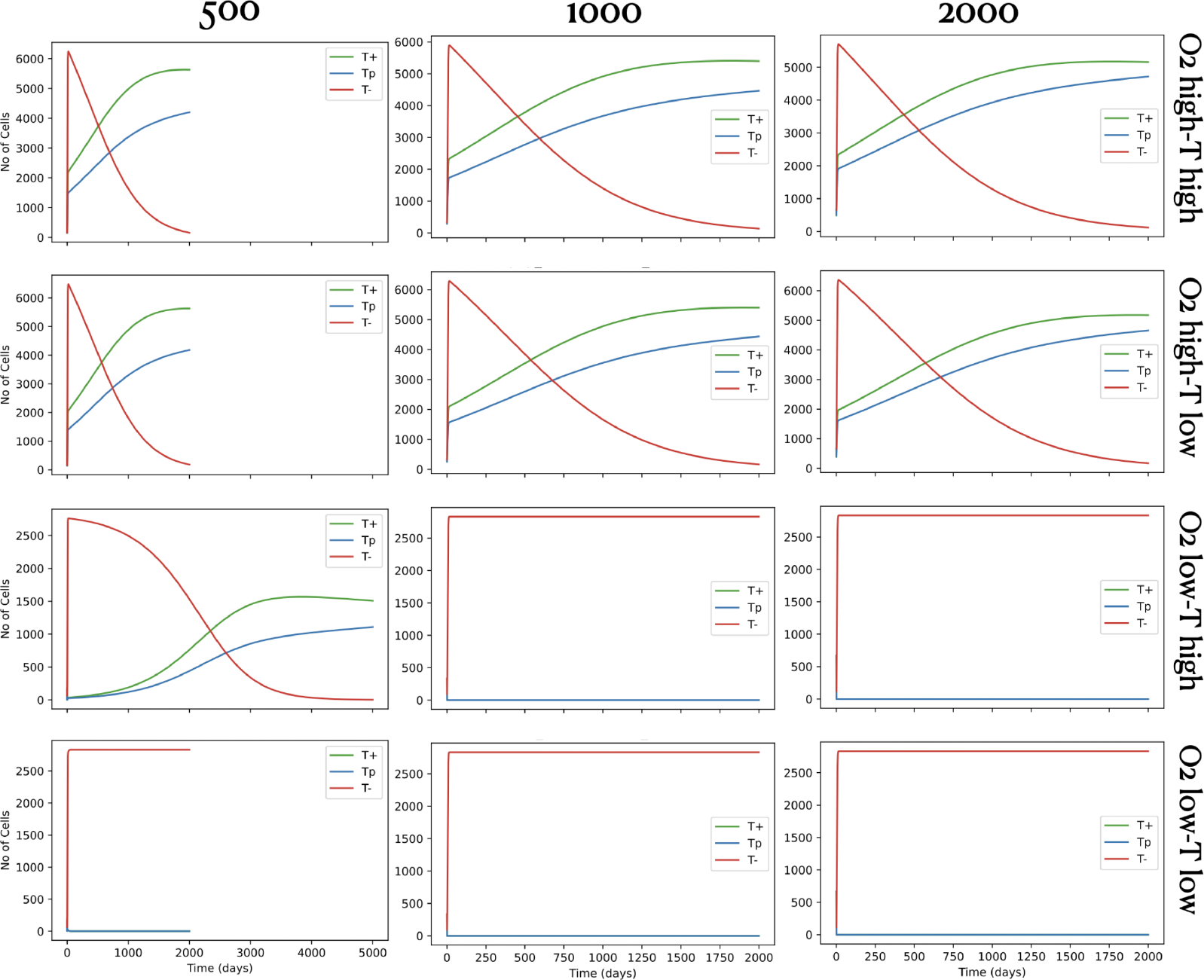
Cell growth time series when <u>only testosterone</u> is supplemented externally. Resource use efficiency combinations vary across rows and initial population size across columns. All simulations were run for 2000 days, except for one case with initial population size 500, low use efficiency of oxygen and high use efficiency of testosterone, which was run for 5000 days for a clear plateau to emerge.

**Supplementary Figure S5:**
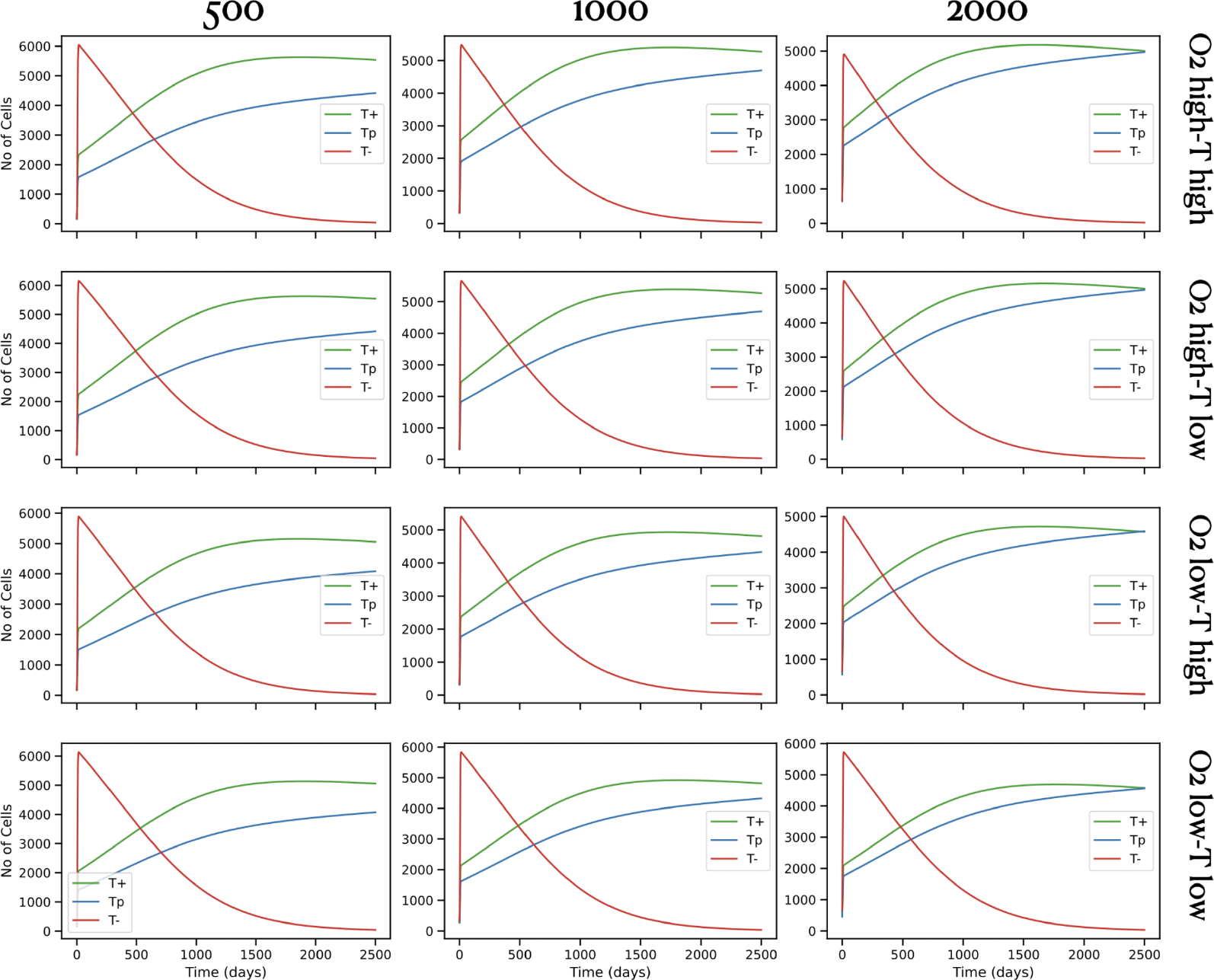
Cell growth time series when <u>both</u> resources are supplemented externally. Resource use efficiency combinations vary across rows and initial population size across columns. These simulations were run for 2500 days.

**Supplementary Figure S6:**
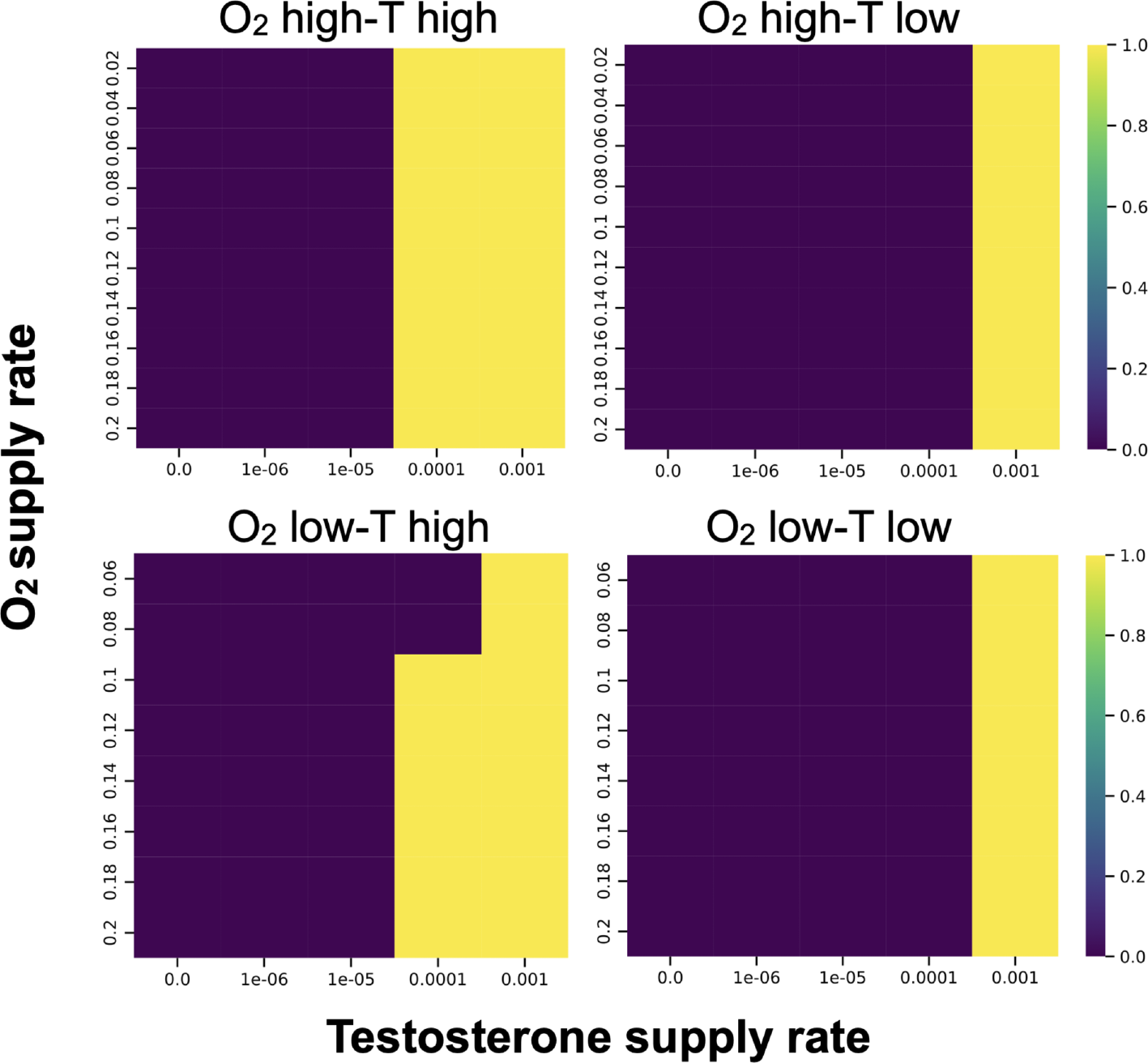
Overall population viability under a range of resource supply conditions. Heat maps of steady state ratio of [*T* ^*p*^ + *T* ^+^] against total population size over a range of supply rates of oxygen and testosterone. By definition, all resource supply states where this ratio goes to zero represents a steady state population of only *T* ^*−*^. Note that under low use efficiency of oxygen, oxygen supply rates below 0.06 min^*−*1^ could not support any cell growth at all and are therefore not shown in this plot. In all cases, simulations were initialised with equal numbers of all three cell types, with an initial population size of 500 and run for 2500 days.

**Supplementary Figure S7:**
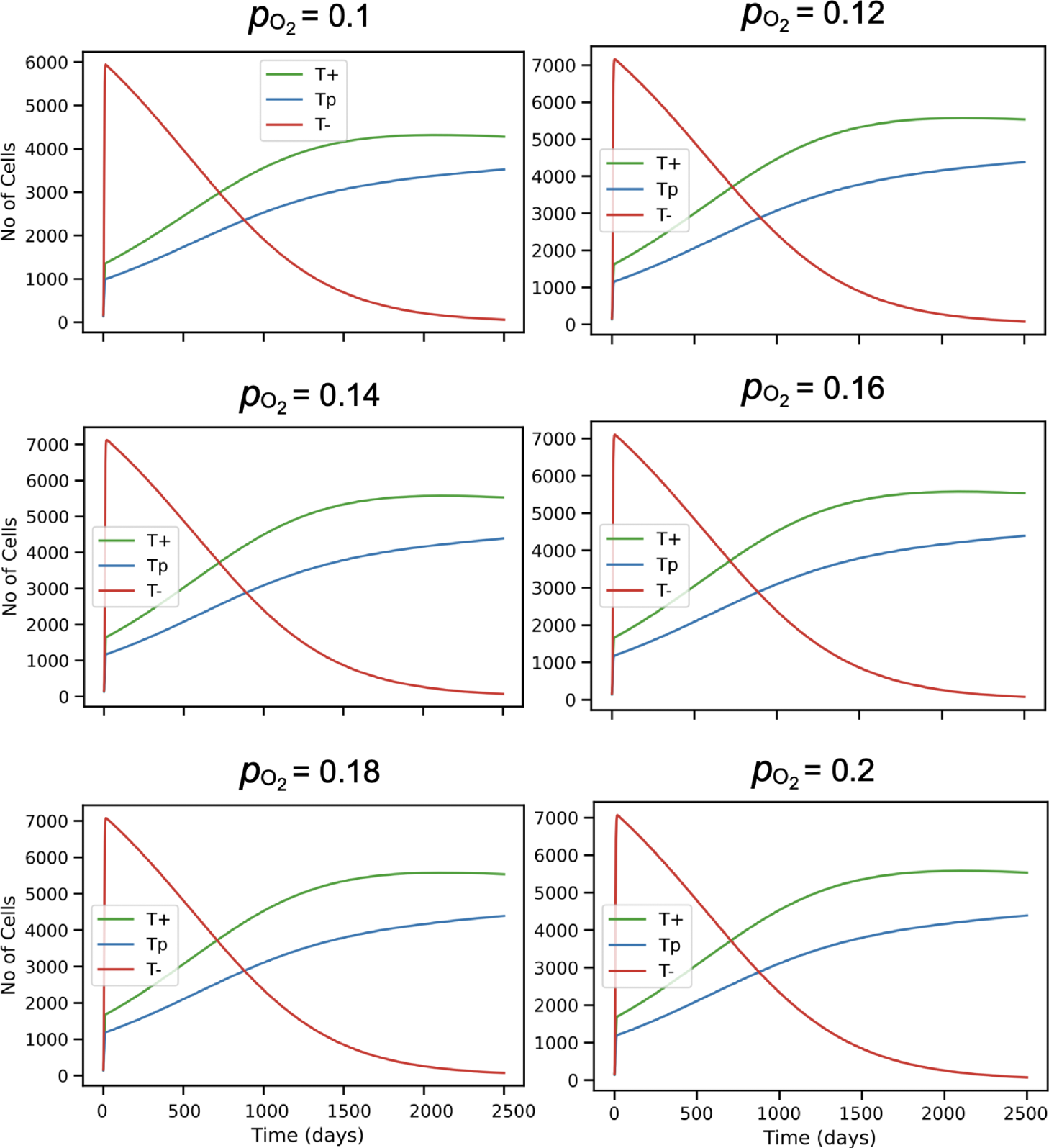
Time series of cell growth corresponding to Figure 4A in the main text for testosterone supply rate 0.0001 min^*−*1^. Low use efficiency of oxygen and high testosterone use efficiency, over a range of oxygen supply rates as specified in the panel titles. Initial population size was 500 with equal numbers of all three cell types.

**Supplementary Figure S8:**
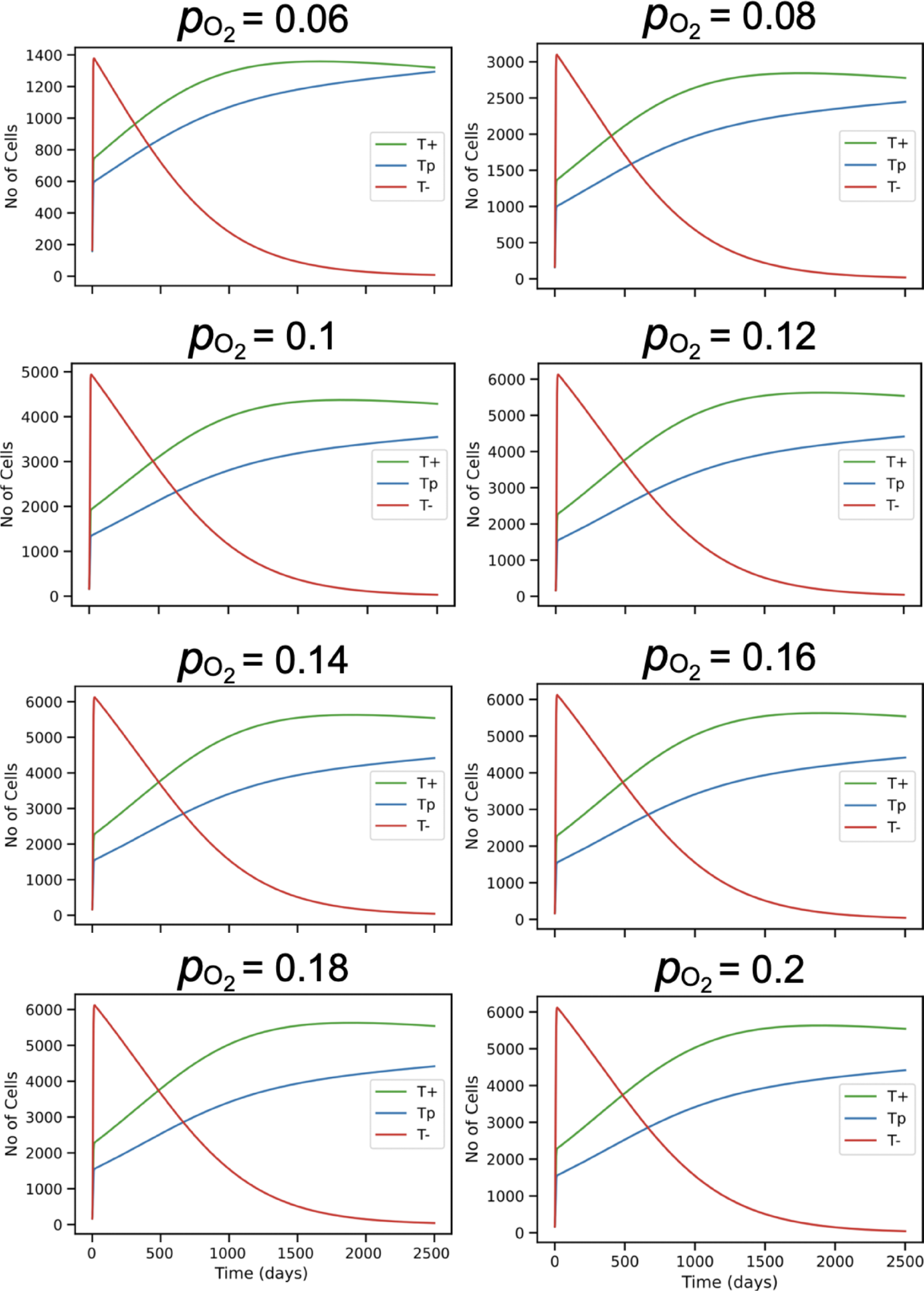
Time series of cell growth corresponding to Figure 4A in the main text for testosterone supply rate 0.001 min^*−*1^. Low use efficiency of oxygen and high testosterone use efficiency, over a range of oxygen supply rates as specified in the panel titles. Initial population size was 500 with equal numbers of all three cell types. Note that cell growth is now viable for lower oxygen supply rates than in the previous figure.

**Supplementary Figure S9:**
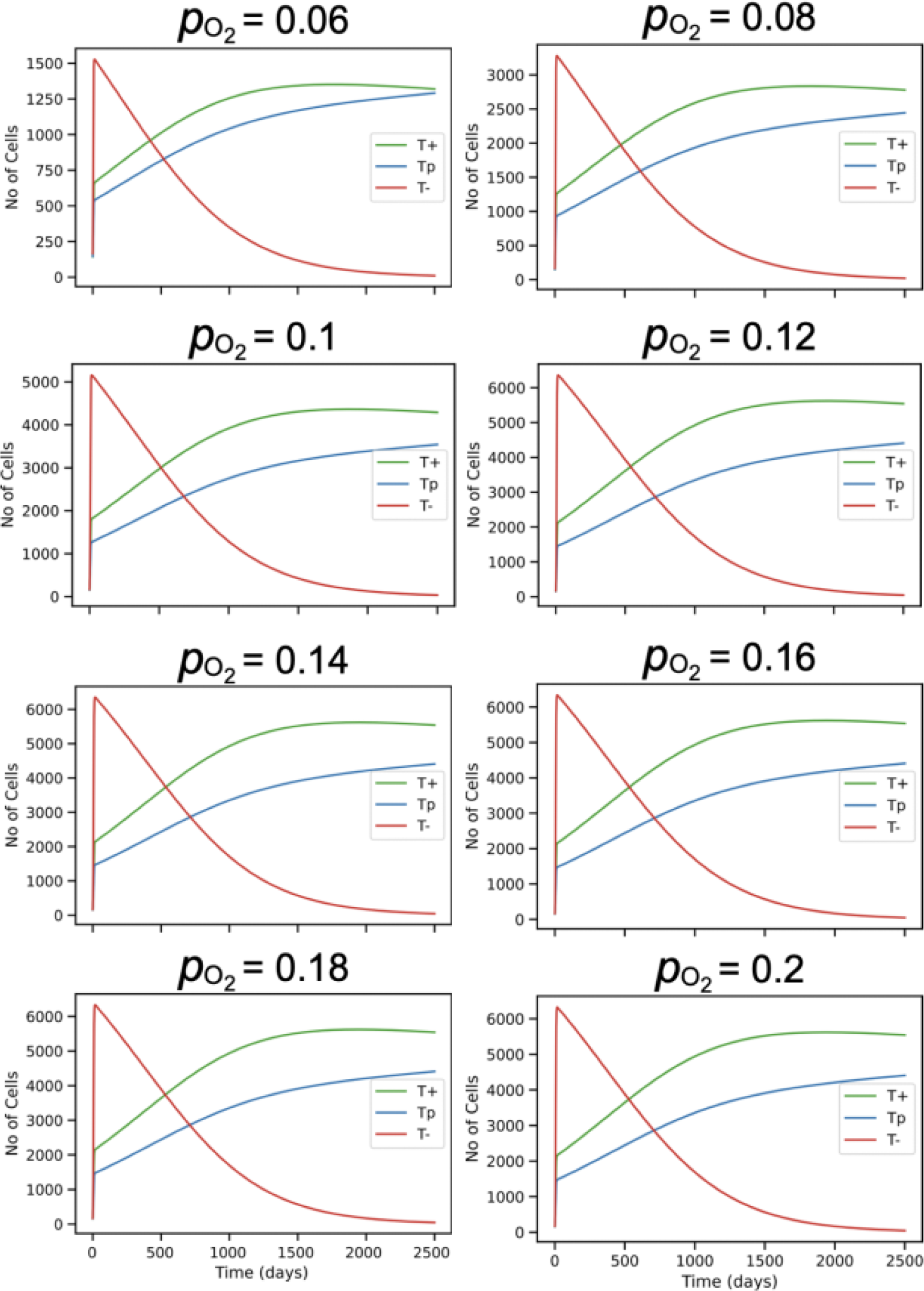
Time series of cell growth corresponding to Figure 4B in the main text. Low use efficiency of both oxygen and testosterone, over a range of oxygen supply rates as specified in the panel titles. Initial population size was 500 with equal numbers of all three cell types, and testosterone supply rate was fixed at 0.001 min^*−*1^.

## References

Marco Archetti, Daniela A Ferraro, and Gerhard Christofori. Heterogeneity for IGF-II production maintained by public goods dynamics in neuroendocrine pancreatic cancer. Proc Natl Acad Sci U S A, 112(6):1–6, 2015. ISSN 1091-6490. doi: 10.1073/pnas.1414653112. arXiv: 1408.1149 ISBN: 1215421109.

Amos Bairoch. The Cellosaurus, a Cell-Line Knowledge Resource. Journal of Biomolecular Techniques : JBT, 29(2):25–38, July 2018. ISSN 1524-0215. doi: 10.7171/jbt.18-2902-002.

D. Basanta, J. G. Scott, M. N. Fishman, G. Ayala, S. W. Hayward, and A. R. A. Anderson. Investigating prostate cancer tumour–stroma interactions: clinical and biological insights from an evolutionary game. British Journal of Cancer, 106(1):174–181, January 2012. ISSN 1532-1827. doi: 10.1038/bjc.2011.517. Number: 1 Publisher: Nature Publishing Group.

Kävan ç Birsoy, Richard Possemato, Franziska K. Lorbeer, Erol C. Bayraktar, Prathapan Thiru, Burcu Yucel, Tim Wang, Walter W. Chen, Clary B. Clish, and David M. Sabatini. Metabolic determinants of cancer cell sensitivity to glucose limitation and biguanides. Nature, 508(1):108–112, April 2014. ISSN 14764687. doi: 10.1038/nature13110. arXiv: NIHMS150003 Publisher: Nature Publishing Group ISBN: 1476-4687 (Electronic) \ r0028-0836 (Linking).

Renee Brady-Nicholls, John D. Nagy, Travis A. Gerke, Tian Zhang, Andrew Z. Wang, Jingsong Zhang, Robert A. Gatenby, and Heiko Enderling. Prostate-specific antigen dynamics predict individual responses to intermittent androgen deprivation. Nature Communications, 11(1):1750, April 2020. ISSN 2041-1723. doi: 10.1038/s41467-020-15424-4. Number: 1 Publisher: Nature Publishing Group.

Louis Calistro Alvarado. Population differences in the testosterone levels of young men are associated with prostate cancer disparities in older men. American Journal of Human Biology, 22(4):449–455, 2010. ISSN 1520-6300. doi: 10.1002/ajhb.21016. eprint: https://onlinelibrary.wiley.com/doi/pdf/10.1002/ajhb.21016.

American Cancer Society. Cancer Facts & Figures 2023. American Cancer Society, 2023.

Suzanne Carreira, Alessandro Romanel, Jane Goodall, Emily Grist, Roberta Ferraldeschi, Susana Miranda, Davide Prandi, David Lorente, Jean-Sebastien Frenel, Carmel Pezaro, Aurelius Omlin, Daniel Nava Rodrigues, Penelope Flohr, Nina Tunariu, Johann S. de Bono, Francesca Demichelis, and Gerhardt Attard. Tumor clone dynamics in lethal prostate cancer. Science Translational Medicine, 6(254):254ra125–254ra125, September 2014. doi: 10.1126/scitranslmed.3009448. Publisher: American Association for the Advancement of Science.

Michelle T Chan, Gareth E Lim, Søs Skovsø, Yu Hsuan Carol Yang, Tobias Albrecht, Emilyn U Alejandro, Corinne A Hoesli, James M Piret, Garth L Warnock, and James D Johnson. Effects of insulin on human pancreatic cancer progression modeled in vitro. BMC Cancer, 14(1):814, December 2014. ISSN 1471-2407. doi: 10.1186/1471-2407-14-814. Publisher: BioMed Central ISBN: 1471-2407 (Electronic)\r1471-2407 (Linking).

Chih-Pin Chuu, John M. Kokontis, Richard A. Hiipakka, Junichi Fukuchi, Hui-Ping Lin, Ching-Yu Lin, Chiech Huo, Liang-Cheng Su, and Shutsung Liao. Androgen suppresses proliferation of castration-resistant LNCaP 104-R2 prostate cancer cells through androgen receptor, Skp2, and c-Myc. Cancer Science, 102(11):2022–2028, 2011. ISSN 1349-7006. doi: 10.1111/j.1349-7006.2011.02043.x. eprint: https://onlinelibrary.wiley.com/doi/pdf/10.1111/j.1349-7006.2011.02043.x.

Z. Culig, J. Hoffmann, M. Erdel, I. E. Eder, A. Hobisch, A. Hittmair, G. Bartsch, G. Utermann, M. R. Schneider, K. Parczyk, and H. Klocker. Switch from antagonist to agonist of the androgen receptor blocker bicalutamide is associated with prostate tumour progression in a new model system. British Journal of Cancer, 81(2):242–251, September 1999. ISSN 1532-1827. doi: 10.1038/sj.bjc.6690684. Number: 2 Publisher: Nature Publishing Group.

Jessica J. Cunningham, Joel S. Brown, Robert A. Gatenby, and Kate ř ina Sta ň ková. Optimal control to develop therapeutic strategies for metastatic castrate resistant prostate cancer. Journal of Theoretical Biology, 459:67–78, December 2018. ISSN 10958541. doi: 10.1016/j.jtbi.2018.09.022. Publisher: Academic Press.

Paulette R. Dillard, Ming-Fong Lin, and Shafiq A. Khan. Androgen-independent prostate cancer cells acquire the complete steroidogenic potential of synthesizing testosterone from cholesterol. Molecular and Cellular Endocrinology, 295(1):115–120, November 2008. ISSN 0303-7207. doi: 10.1016/j.mce.2008.08.013.

Nathan Farrokhian, Jeff Maltas, Mina Dinh, Arda Durmaz, Patrick Ellsworth, Masahiro Hitomi, Erin McClure, Andriy Marusyk, Artem Kaznatcheev, and Jacob G. Scott. Measuring competitive exclusion in non–small cell lung cancer. Science Advances, 8(26):eabm7212, July 2022. doi: 10.1126/sciadv.abm7212. Publisher: American Association for the Advancement of Science.

Duc Fehr, Harini Veeraraghavan, Andreas Wibmer, Tatsuo Gondo, Kazuhiro Matsumoto, Herbert Alberto Vargas, Evis Sala, Hedvig Hricak, and Joseph O. Deasy. Automatic classification of prostate cancer Gleason scores from multiparametric magnetic resonance images. Proceedings of the National Academy of Sciences, 112(46):E6265–E6273, 2015. doi: 10.1073/pnas.1505935112.

Fabrizio Fontana, Martina Anselmi, and Patrizia Limonta. Molecular mechanisms and genetic alterations in prostate cancer: From diagnosis to targeted therapy. Cancer Letters, 534:215619, May 2022. ISSN 0304-3835. doi: 10.1016/j.canlet.2022.215619.

Jill A. Gallaher, Pedro M. Enriquez-Navas, Kimberly A. Luddy, Robert A. Gatenby, and Alexander R.A. Anderson. Spatial Heterogeneity and Evolutionary Dynamics Modulate Time to Recurrence in Continuous and Adaptive Cancer Therapies. Cancer Research, 78(8):2127–2139, April 2018. ISSN 0008-5472. doi: 10.1158/0008-5472.CAN-17-2649. Publisher: American Association for Cancer Research.

Robert A. Gatenby, Ariosto S. Silva, Robert J. Gillies, and B. Roy Frieden. Adaptive therapy. Cancer Research, 69(11):4894–4903, 2009. ISSN 00085472. doi: 10.1158/0008-5472.CAN-08-3658.

Craig Gedye and Vishal Navani. Find the path of least resistance: Adaptive therapy to delay treatment failure and improve outcomes. Biochimica et Biophysica Acta (BBA) - Reviews on Cancer, 1877(2):188681, March 2022. ISSN 0304-419X. doi: 10.1016/J.BBCAN.2022.188681. Publisher: Elsevier.

Ahmadreza Ghaffarizadeh, Randy Heiland, Samuel H. Friedman, Shannon M. Mumenthaler, and Paul Macklin. PhysiCell: An open source physics-based cell simulator for 3-D multicellular systems. PLoS Computational Biology, 14(2):e1005991, February 2018. ISSN 15537358. doi: 10.1371/journal.pcbi.1005991. Publisher: Public Library of Science.

Robert J. Gillies, Paul E. Kinahan, and Hedvig Hricak. Radiomics: Images are more than pictures, they are data. Radiology, 278(2):563–577, February 2016. ISSN 0033-8419. doi: 10.1148/radiol.2015151169. Publisher: Radiological Society of North America.

Jennifer Gordetsky and Jonathan Epstein. Grading of prostatic adenocarcinoma: current state and prognostic implications. Diagnostic Pathology, 11(1):25, March 2016. ISSN 1746-1596. doi: 10.1186/s13000-016-0478-2.

Catherine S. Grasso, Yi-Mi Wu, Dan R. Robinson, Xuhong Cao, Saravana M. Dhanasekaran, Amjad P. Khan, Michael J. Quist, Xiaojun Jing, Robert J. Lonigro, J. Chad Brenner, Irfan A. Asangani, Bushra Ateeq, Sang Y. Chun, Javed Siddiqui, Lee Sam, Matt Anstett, Rohit Mehra, John R. Prensner, Nallasivam Palanisamy, Gregory A. Ryslik, Fabio Vandin, Benjamin J. Raphael, Lakshmi P. Kunju, Daniel R. Rhodes, Kenneth J. Pienta, Arul M. Chinnaiyan, and Scott A. Tomlins. The mutational landscape of lethal castration-resistant prostate cancer. Nature, 487(7406):239–243, July 2012. ISSN 1476-4687. doi: 10.1038/nature11125. Number: 7406 Publisher: Nature Publishing Group.

C. W. Gregory, R. T. Johnson, J. L. Mohler, F. S. French, and E. M. Wilson. Androgen receptor stabilization in recurrent prostate cancer is associated with hypersensitivity to low androgen. Cancer Research, 61(7):2892–2898, April 2001. ISSN 0008-5472.

James P. Grover. Resource Competition. Springer Science & Business Media, July 1997. ISBN 978-0-412-74930-8. Google-Books-ID: x89Rl1nYR8gC.

Numsen Hail, Ping Chen, and Lane R. Bushman. Teriflunomide (Leflunomide) Promotes Cytostatic, Antioxidant, and Apoptotic Effects in Transformed Prostate Epithelial Cells: Evidence Supporting a Role for Teriflunomide in Prostate Cancer Chemoprevention. Neoplasia, 12(6):464–475, June 2010. ISSN 1476-5586. doi: 10.1593/neo.10168.

Douglas Hanahan and Robert A. Weinberg. Hallmarks of Cancer: The Next Generation. Cell, 144(5):646–674, 2011. ISSN 00928674. doi: 10.1016/j.cell.2011.02.013.

Elsa Hansen and Andrew F. Read. Cancer therapy: Attempt cure or manage drug resistance? Evolutionary Applications, 13(7):1660–1672, August 2020. ISSN 1752-4571. doi: 10.1111/eva.12994. Publisher: Wiley.

Elsa Hansen, Robert J. Woods, and Andrew F. Read. How to Use a Chemotherapeutic Agent When Resistance to It Threatens the Patient. PLOS Biology, 15(2):e2001110, February 2017. ISSN 1545-7885. doi: 10.1371/journal.pbio.2001110. Publisher: Public Library of Science.

Harsh Vardhan Jain, Steven K Clinton, Arvinder Bhinder, and Avner Friedman. Mathematical modeling of prostate cancer progression in response to androgen ablation therapy. Proceedings of the National Academy of Sciences, 108(49):19701–19706, December 2011. ISSN 0027-8424. doi: 10.1073/pnas.1115750108. Publisher: National Academy of Sciences ISBN: 1091-6490 (Electronic)\r0027-8424 (Linking).

Irina Kareva and Joel S. Brown. Estrogen as an Essential Resource and the Coexistence of ER+ and ER– Cancer Cells. Frontiers in Ecology and Evolution, 9, 2021. ISSN 2296-701X.

Irina Kareva, Benjamin Morin, and Carlos Castillo-Chavez. Resource Consumption, Sustainability, and Cancer. Bulletin of Mathematical Biology, 77(2):319–338, February 2015. ISSN 15229602. doi: 10.1007/S11538-014-9983-1/FIGURES/4. Publisher: Springer Science and Business Media, LLC.

Sara Loponte, Sara Lovisa, Angela K. Deem, Alessandro Carugo, and Andrea Viale. The Many Facets of Tumor Heterogeneity: Is Metabolism Lagging Behind? Cancers, 11 (10):1574, October 2019. ISSN 2072-6694. doi: 10.3390/cancers11101574. Number: 10 Publisher: Multidisciplinary Digital Publishing Institute.

Esha Madan, Ant ó nio M. Palma, Vignesh Vudatha, Jose G. Trevino, Kedar Nath Natarajan, Robert A. Winn, Kyoung Jae Won, Trevor A. Graham, Ronny Drapkin, Stuart A.C. McDonald, Paul B. Fisher, and Rajan Gogna. Cell Competition in Carcinogenesis. Cancer Research, 82(24):4487–4496, December 2022. ISSN 0008-5472. doi: 10.1158/0008-5472.CAN-22-2217.

Urko M Marigorta, Lee A Denson, Jeffrey S Hyams, Kajari Mondal, Jarod Prince, Thomas D Walters, Anne Griffiths, Joshua D Noe, Wallace V Crandall, Joel R Rosh, David R Mack, Richard Kellermayer, Melvin B Heyman, Susan S Baker, Michael C Stephens, Robert N Baldassano, James F Markowitz, Mi-Ok Kim, Marla C Dubinsky, Judy Cho, Bruce J Aronow, Subra Kugathasan, and Greg Gibson. Transcriptional risk scores link GWAS to eQTLs and predict complications in Crohn’s disease. Nature Genetics, August 2017. ISSN 1061-4036. doi: 10.1038/ng.3936.

S D Martin, G Coukos, R A Holt, and B H Nelson. Targeting the undruggable: immunotherapy meets personalized oncology in the genomic era. Annals of oncology : official journal of the European Society for Medical Oncology / ESMO, 26(12):2367–74, December 2015. ISSN 1569-8041. doi: 10.1093/annonc/mdv382.

Andriy Marusyk, Doris P. Tabassum, Philipp M. Altrock, Vanessa Almendro, Franziska Michor, and Kornelia Polyak. Non-cell-autonomous driving of tumour growth supports sub-clonal heterogeneity. Nature, 514(7520):54–58, October 2014. ISSN 1476-4687. doi: 10.1038/nature13556. Number: 7520 Publisher: Nature Publishing Group.

James L. Mohler, Christopher W. Gregory, O. Harris Ford, III, Desok Kim, Catharina M. Weaver, Peter Petrusz, Elizabeth M. Wilson, and Frank S. French. The Androgen Axis in Recurrent Prostate Cancer. Clinical Cancer Research, 10(2):440–448, February 2004. ISSN 1078-0432. doi: 10.1158/1078-0432.CCR-1146-03.

R. Bruce Montgomery, Elahe A. Mostaghel, Robert Vessella, David L. Hess, Thomas F. Kalhorn, Celestia S. Higano, Lawrence D. True, and Peter S. Nelson. Maintenance of Intratumoral Androgens in Metastatic Prostate Cancer: A Mechanism for Castration-Resistant Tumor Growth. Cancer Research, 68(11):4447–4454, June 2008. ISSN 0008-5472. doi: 10.1158/0008-5472.CAN-08-0249.

Rodolfo Montironi, Alessia Cimadamore, Liang Cheng, Antonio Lopez-Beltran, and Marina Scarpelli. Prostate cancer grading in 2018: limitations, implementations, cribriform morphology, and biological markers. The International Journal of Biological Markers, 33(4):331–334, November 2018. ISSN 0393-6155. doi: 10.1177/1724600818781296. Publisher: SAGE Publications Ltd STM.

Mario E. Muscarella and James P. O’Dwyer. Species dynamics and interactions via metabolically informed consumer-resource models. Theoretical Ecology, 13(4):503–518, December 2020. ISSN 1874-1746. doi: 10.1007/s12080-020-00466-7.

Saroj Niraula, Arnoud J. Templeton, Francisco E. Vera-Badillo, Anthony M. Joshua, Srikala S. Sridhar, Peter W. Cheung, Paul M. Yip, Anna Dodd, Zoann Nugent, and Ian F. Tannock. Study of testosterone-guided androgen deprivation therapy in management of prostate cancer: Study of Testosterone-Guided Androgen Deprivation. The Prostate, 76 (2):235–242, February 2016. ISSN 02704137. doi: 10.1002/pros.23117.

Tsutomu Nishiyama and Tatsuhiko Hoshii. Testosterone-guided ADT for prostate cancer. Nature Reviews Urology, 13(4):189–191, April 2016. ISSN 1759-4812, 1759-4820. doi: 10.1038/nrurol.2016.9.

C Kent Osborne, Kim Hobbs, and Gary M Clark. Effect of Estrogens and Antiestrogens on Growth of Human Breast Cancer Cells in Athymic Nude Mice1. CANCER RESEARCH, 45:584–590, 1985.

Stephanie T. Page, Daniel W. Lin, Elahe A. Mostaghel, David L. Hess, Lawrence D. True, John K. Amory, Peter S. Nelson, Alvin M. Matsumoto, and William J. Bremner. Persistent Intraprostatic Androgen Concentrations after Medical Castration in Healthy Men. The Journal of Clinical Endocrinology & Metabolism, 91(10):3850–3856, October 2006. ISSN 0021-972X. doi: 10.1210/jc.2006-0968.

Sarthak Sahoo, Ashutosh Mishra, Harsimran Kaur, Kishore Hari, Srinath Muralidharan, Susmita Mandal, and Mohit Kumar Jolly. A mechanistic model captures the emergence and implications of non-genetic heterogeneity and reversible drug resistance in ER+ breast cancer cells. NAR Cancer, 3(3):zcab027, September 2021. ISSN 2632-8674. doi: 10.1093/narcan/zcab027.

Michael Stanbrough, Glenn J. Bubley, Kenneth Ross, Todd R. Golub, Mark A. Rubin, Trevor M. Penning, Phillip G. Febbo, and Steven P. Balk. Increased Expression of Genes Converting Adrenal Androgens to Testosterone in Androgen-Independent Prostate Cancer. Cancer Research, 66(5):2815–2825, March 2006. ISSN 0008-5472. doi: 10.1158/0008-5472.CAN-05-4000.

Grant D. Stewart, James A. Ross, Duncan B. McLaren, Christopher C. Parker, Fouad K. Habib, and Antony C.P. Riddick. The relevance of a hypoxic tumour microenvironment in prostate cancer. BJU International, 105(1):8–13, 2010. ISSN 1464-410X. doi: 10.1111/j.1464-410X.2009.08921.x. eprint: https://onlinelibrary.wiley.com/doi/pdf/10.1111/j.1464-410X.2009.08921.x.

Benjamin A. Teply, Hao Wang, Brandon Luber, Rana Sullivan, Irina Rifkind, Ashley Bruns, Avery Spitz, Morgan DeCarli, Victoria Sinibaldi, Caroline F. Pratz, Changxue Lu, John L. Silberstein, Jun Luo, Michael T. Schweizer, Charles G. Drake, Michael A. Carducci, Channing J. Paller, Emmanuel S. Antonarakis, Mario A. Eisenberger, and Samuel R. Denmeade. Bipolar androgen therapy in men with metastatic castration-resistant prostate cancer after progression on enzalutamide: an open-label, phase 2, multicohort study. The Lancet Oncology, 19(1):76–86, January 2018. ISSN 14745488. doi: 10.1016/S1470-2045(17)30906-3. Publisher: Lancet Publishing Group.

David Tilman. Resources: A Graphical-Mechanistic Approach to Competition and Predation. The American Naturalist, 116(3):362–393, September 1980. ISSN 0003-0147. doi: 10.1086/283633. Publisher: The University of Chicago Press.

Mark A. Titus, Michael J. Schell, Fred B. Lih, Kenneth B. Tomer, and James L. Mohler. Testosterone and Dihydrotestosterone Tissue Levels in Recurrent Prostate Cancer. Clinical Cancer Research, 11(13):4653–4657, July 2005. ISSN 1078-0432. doi: 10.1158/1078-0432.CCR-05-0525.

Y Umekita, R A Hiipakka, J M Kokontis, and S Liao. Human prostate tumor growth in athymic mice: inhibition by androgens and stimulation by finasteride. Proceedings of the National Academy of Sciences, 93(21):11802–11807, October 1996. doi: 10.1073/pnas.93.21.11802. Publisher: Proceedings of the National Academy of Sciences.

Philip A. Watson, Vivek K. Arora, and Charles L. Sawyers. Emerging mechanisms of resistance to androgen receptor inhibitors in prostate cancer. Nature Reviews Cancer, 15(12): 701–711, December 2015. ISSN 1474-1768. doi: 10.1038/nrc4016. Number: 12 Publisher: Nature Publishing Group.

Jeffrey West, Li You, Jingsong Zhang, Robert A. Gatenby, Joel S. Brown, Paul K. Newton, and Alexander R.A. Anderson. Towards Multidrug Adaptive Therapy. Cancer Research, 80(7):1578–1589, April 2020. ISSN 0008-5472. doi: 10.1158/0008-5472.CAN-19-2669.

Andreas Wibmer, Hedvig Hricak, Tatsuo Gondo, Kazuhiro Matsumoto, Harini Veeraraghavan, Duc Fehr, Junting Zheng, Debra Goldman, Chaya Moskowitz, Samson W. Fine, Victor E. Reuter, James Eastham, Evis Sala, and Hebert Alberto Vargas. Haralick texture analysis of prostate MRI: utility for differentiating non-cancerous prostate from prostate cancer and differentiating prostate cancers with different Gleason scores. European Radiology, 25(10):2840–2850, October 2015. ISSN 1432-1084. doi: 10.1007/s00330-015-3701-8.

Jingsong Zhang, Jessica J. Cunningham, Joel S. Brown, and Robert A. Gatenby. Integrating evolutionary dynamics into treatment of metastatic castrate-resistant prostate cancer. Nature Communications, 8(1):1816, December 2017. ISSN 20411723. doi: 10.1038/s41467-017-01968-5. Publisher: Nature Publishing Group.

Jingsong Zhang, Jessica Cunningham, Joel Brown, and Robert Gatenby. Evolution-based mathematical models significantly prolong response to abiraterone in metastatic castrate-resistant prostate cancer and identify strategies to further improve outcomes. eLife, 11: e76284, June 2022. ISSN 2050-084X. doi: 10.7554/eLife.76284.

Jie Zheng. Energy metabolism of cancer: Glycolysis versus oxidative phosphorylation (review). Oncology Letters, 4(6):1151–1157, December 2012. ISSN 17921074. doi: 10.3892/ol.2012.928. arXiv: NIHMS150003 Publisher: Spandidos Publications ISBN: 1792-1074\r1792-1082.

## Supplementary References

Thomas A. Stamey, Norman Yang, Alan R. Hay, John E. McNeal, Fuad S. Freiha, and Elise Redwine. Prostate-Specific Antigen as a Serum Marker for Adenocarcinoma of the Prostate. New England Journal of Medicine, 317(15):909–916, October 1987. ISSN 0028-4793, 1533-4406. doi: 10.1056/NEJM198710083171501.

